# A Modular Platform for Streamlining Automated Cryo-FIB Workflows

**DOI:** 10.1101/2021.05.19.444745

**Authors:** Sven Klumpe, Herman K. H. Fung, Sara K. Goetz, Ievgeniia Zagoriy, Bernhard Hampoelz, Xiaojie Zhang, Philipp S. Erdmann, Janina Baumbach, Christoph W. Müller, Martin Beck, Jürgen M. Plitzko, Julia Mahamid

## Abstract

Lamella micromachining by focused ion beam milling at cryogenic temperature (cryo-FIB) has matured into a preparation method widely used for cellular cryo-electron tomography. Due to the limited ablation rates of low Ga^+^ ion beam currents required to maintain the structural integrity of vitreous specimens, current preparation protocols are time-consuming and labor intensive. The improved stability of new generation cryo-FIB instruments now enables automated operations. Here, we present an open-source software tool, SerialFIB, for creating automated and customizable cryo-FIB preparation protocols. The software encompasses a graphical user interface for easy execution of routine lamellae preparations, a scripting module compatible with available python packages, and interfaces with 3-dimensional correlative light and electron microscopy (CLEM) tools. The software enables the streamlining of advanced cryo-FIB protocols such as multi-modal imaging, CLEM guided preparation and *in situ* lamella lift-out procedures. Our software therefore provides a foundation for further development of advanced cryogenic imaging and sample preparation protocols.

## Introduction

Cryo-electron tomography (cryo-ET) is a structural biology technique that can reveal the sociology of macromolecules in their native environment (1). Fixation through rapid cooling to cryogenic temperature (below −140 °C (2, 3)) arrests water molecules in a glass-like phase, thus preserving biological structures in a near-native state. The vitrification temperature is pressure dependent. Hence, the maximum achievable vitrification depth ranges from ~10 μm for plunge-freezing at ambient pressure, to ~ 200 μm in high pressure freezing (HPF) (4). Most cellular cryo-ET imaging is performed at 300 kV, at which the inelastic mean free path of an electron in biological material is in the order of 300–400 nm. Consequently, thicker samples will exhibit a rapidly decreasing signal-to-noise ratio (5–7). Therefore, most cells, excluding the thinnest peripheries of adherent eukaryotic cells, are not directly suitable for cryo-ET. Cryo-FIB micromachining enables the production of electron-transparent cellular slices thinner than 300 nm, which are termed lamellae (8, 9). Cryo-FIB milling has been successfully applied to single-cell specimens deposited or grown directly on transmission electron microscopy (TEM) grids, or to voluminous HPF multicellular samples using *in situ* lift-out (10, 11). These approaches have been shown to yield specimens of suitable quality for cryo-TEM imaging (12), circumventing artifacts described for mechanical sectioning by cryo-ultramicrotomy (13). However, common cryo-FIB preparations constitute a low throughput method owing to slow ablation rates of Ga^+^ ions at low currents (30 pA – 1 nA) typically used to micromachine vitreous specimens. Furthermore, it requires significant user expertise with a steep learning curve, which limits its practical usability. In addition, emerging concepts such as multi-modal imaging through cryo-FIB-scanning electron microscopy (SEM) volume imaging coupled to lamella preparation, are challenging to achieve with manual operations (14–16). Automation is thus essential to improve the performance, throughput, and applicability of such time-consuming approaches.

Considering the advance in cryo-EM imaging owing to the availability of open source, high-level automation interfaces (17–19), cryo-FIB-SEM sample preparation and imaging should similarly benefit from modular automation platforms. Automation for cryo-FIB lamella preparation has been recently presented, based either on proprietary software (20–22) or on command-line operation (23). Those workflows were designed for the specialized task of preparing routine on-grid lamellae of single-cell specimens. Thus, an open-source and easy-to-use package for customized cryo-FIB-SEM workflows is currently missing. Inspired by the robustness and architecture of SerialEM (24), we developed a user-friendly and customizable automation software platform for cryo-FIB instruments. We demonstrate its flexibility on several use cases ranging from standard preparations to specialized applications. This involves (i) on-grid lamella preparation, (ii) cryo-fluorescence light microscopy (cryo-FLM) based 3D-targeted lamella preparation, (iii) cryo-FIB-SEM volume imaging, with an option for subsequent lamella preparation, and (iv) micromachining for *in situ* lift-out from voluminous HPF samples. Apart from the proprietary application programming interface (API) required for communication with the microscope, the software code is open-source, paving the way for further workflow development. We show that our milling protocols can be performed without supervision on six different specimens. Some of the specimens contained high-density intracellular structures, i.e. lipid droplets or minerals, that introduce additional difficulty to the generation of homogenously thin lamellae. We therefore provide milling recipes optimized on a broad range of cellular samples. In addition, user-tailored time-consuming cryo-FIB-SEM volume imaging and lift-out trench milling can be performed overnight given compatibility of the instrument for prolonged operation times, thereby increasing throughput for advanced imaging and sample preparation.

## Results

### Software Design

In developing SerialFIB, we seek to provide a flexible platform for the creation and execution of customized cryo-FIB milling protocols (Figure 1). The graphical user interface (GUI) and image processing steps are therefore decoupled from underlying communications with the cryo-FIB-SEM instrument. Communication with the instrument is handled through an intermediary “driver”, which makes use of the microscope’s API to perform beam, stage and imaging operations (Figure 1). Here, we developed a driver for Thermo Fisher Scientific (TFS) systems based on the proprietary API AutoScript 4. Automated cryo-FIB workflows were tested on a TFS Scios and two Aquilos systems.

**Figure 1:**
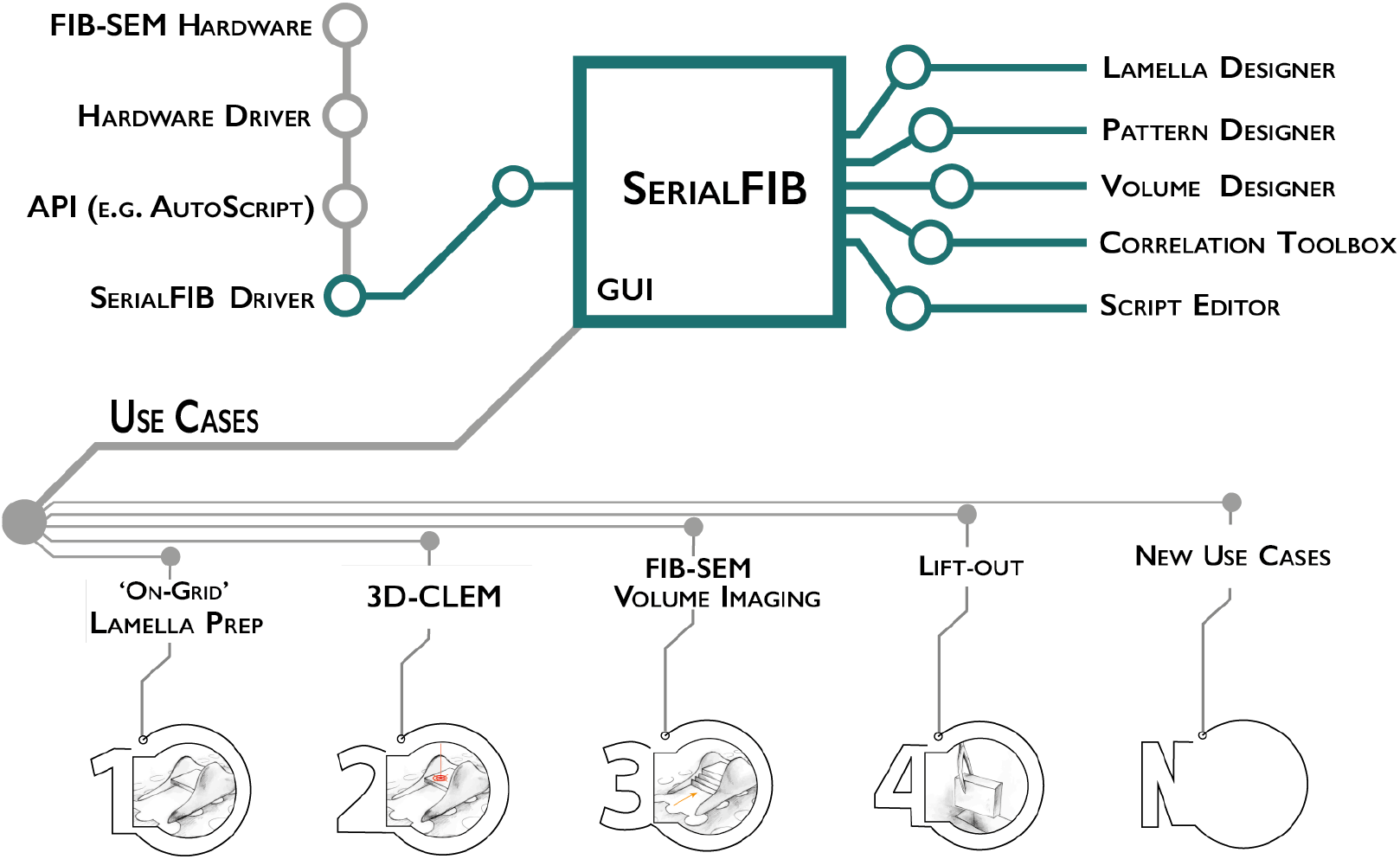
Software architecture, modules and use cases of SerialFIB. Developments presented in this work are highlighted in green. The graphical user interface (GUI) is largely decoupled from instrument operations, which are controlled by the developed SerialFIB Driver. Modules (right) enable design of protocols for different use cases (bottom, 1-N), and a scripting interface that offers flexibility for new developments.

The basic unit for milling or patterning tasks in SerialFIB are “pattern sequence files’’, wherein a sequence of milling operations characterized by the milling current, milling time, pattern size, and pattern position relative to a per-position user-defined reference point, are defined. We provide “protocols”, text-based files that can be translated by SerialFIB into pattern sequence files, defining the sequence of patterning steps to be performed, for a number of use cases. These simplify the creation of pattern sequence files by focusing on use-case-specific parameters, e.g., lamella and slice thickness for on-grid lamella preparation and volume imaging applications, respectively. The use cases include on-grid lamella preparation, 3D cryo-FLM-guided lamella preparation, and FIB-SEM volume imaging. More specialized tasks employ pattern sequence files directly, exemplified here by the lift-out site preparation. Upon selection of a protocol in the SerialFIB GUI (Figure 1-figure supplement 1A), the software automatically generates the necessary pattern sequence files, which are subsequently parsed and executed through the system-specific intermediary driver. Protocol and pattern sequence files can be edited in a text-editor or through their respective “Designer” module, which provides a graphical interface and parameter preview to enable adjustments during parameter optimization (Figure 1-figure supplement 1B-C).

During a typical cryo-FIB session, points of interest on the sample and their coincidence height (the height at which the FIB and SEM beam coincide) are first identified manually through the microscope user interface. Then, in SerialFIB, positions are stored, reference images for realignment are taken, and target position and milling geometry are defined on the reference image per point of interest. For on-grid lamella preparations, for example, the lamella position, width, and extreme points above and below the target lamella from which ablation of material will begin are defined (Figure 1-figure supplement 1A). To start the procedure, the corresponding protocol is initiated. The setup within the SerialFIB GUI is standardized, thus allowing several protocols to be executed at the same position, e.g., FIB-SEM volume imaging (Figure 1-figure supplement 2A) and subsequent lamella preparation (Figure 1-figure supplement 1B).

FIB milling routines that are not covered by our predefined protocols (e.g. Waffle method (25)) can be developed through customized pattern sequence files or the ScriptEditor (Figure 1-figure supplement 2B). In the ScriptEditor, images and stage positions previously defined in the GUI are accessible together with the underlying python commands of the driver. Thus, additional use cases based on custom python scripts can be implemented and shared.

### Automated on-grid lamellae preparation

Similar to recent developments (20, 21, 23), SerialFIB offers an automated solution for preparing on-grid lamellae for cryo-ET. For each position of interest, the user defines the target lamella width and position, as well as extreme milling points on the reference image (Supplementary video 1). The definition of extreme milling points ensures that the milling of grid bars and other objects, which can deflect the ion beam and cause damage to the lamella, is avoided (Figure 2A). Then, in an automated fashion based on the chosen protocols, micro-expansion joints (26) are created to relieve tension in the frozen specimen (Figure 2B), followed by removal of material in two separate stages: (i) rough milling, where all positions of interest are milled to ~1 μm thickness, and (ii) polishing, where the generated slices are thinned to ~200 nm thickness to achieve electron transparency (Figure 2C). By polishing all positions within a short time window, amorphous ice condensation on the lamella is minimized (27).

**Figure 2:**
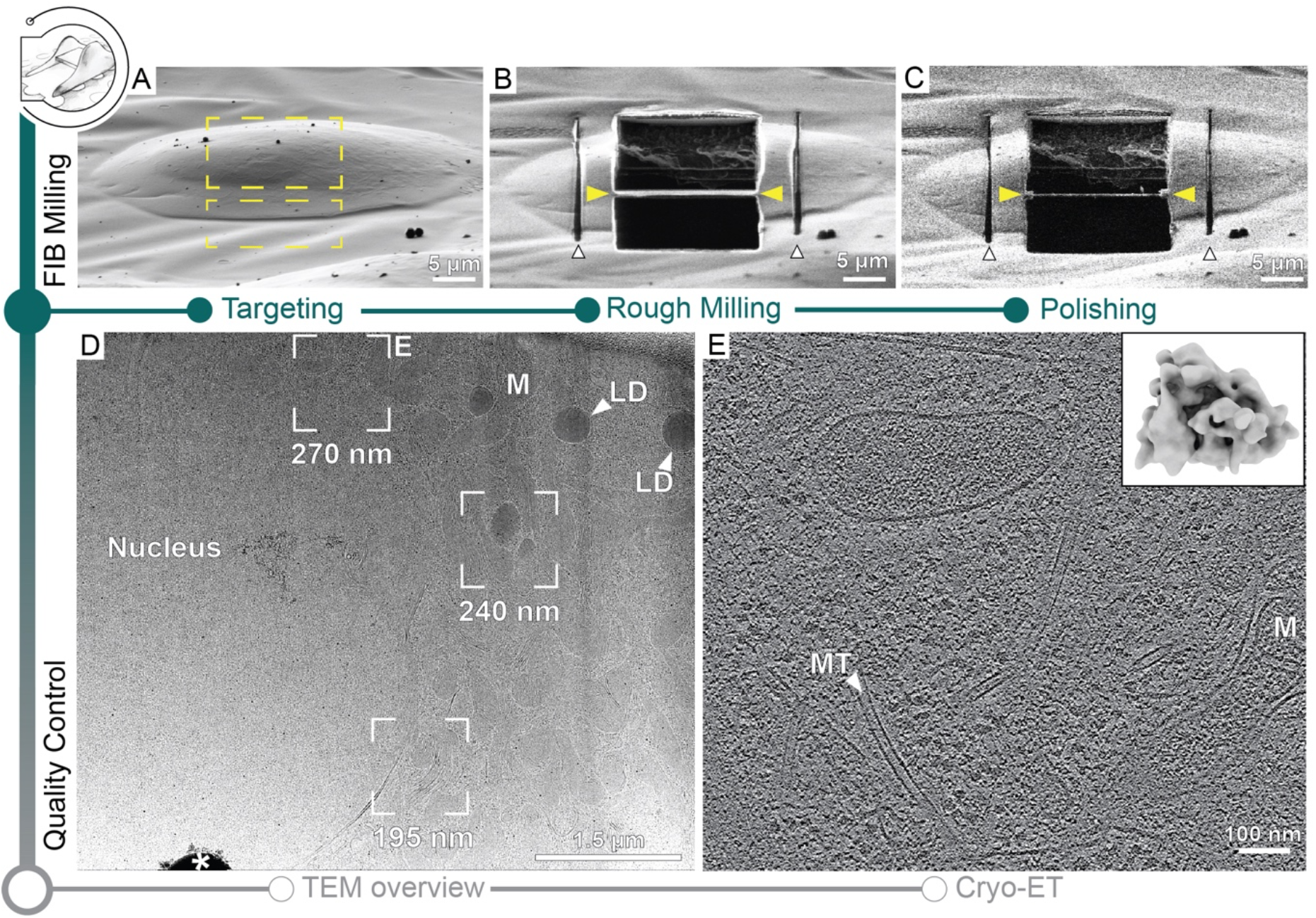
Automated on-grid lamella preparation of Sum159 breast cancer cells. (A) FIB image of a cell prior to lamella preparation. Yellow rectangles indicate milling patterns where material is subsequently removed. (B) Micro-expansion joints (white arrowheads) and lamella (yellow arrowheads) milled to a target thickness of 1 μm. (C) Lamella after polishing to a target thickness of 300 nm. (D) TEM overview of lamella in C. Frames indicate examples of tilt-series acquisition positions (out of 8 acquired on this lamella) and the local thickness determined from reconstructed tomograms. Nucleus, lipid droplets (LD) and mitochondria (M) are observable. * indicates an ice crystal introduced during transfer between FIB and TEM. Area indicated by E is enlarged. (E) A slice through the tomogram depicts the cytosol with microtubules (MT) and a mitochondrion (M). Inset shows a ribosome subtomogram average determined from the dataset collected on this single lamella (4 tomograms; 4378 subtomograms; 24 Å resolution).

Automated milling in SerialFIB uses a series of realignment steps to precisely recall the target position for each milling step. The realignment procedure is based on image cross-correlation (28). It is performed when navigating between stored stage positions and when changing between FIB currents to compensate for imperfect microscope alignments. In the first case, stage movements are used for large distance offsets (> 10 μm) and beam shifts are applied for subtle corrections. A low ion beam current (10 pA) is used to capture the current view for registration with the stored reference image. When changing between FIB currents, two images are taken, one at the previous beam current to serve as a reference, and one at the new beam current to determine the offset. Here, only beam shifts are used for realignment.

For lamella preparation, each milling step is linked to a specific milling time and current. These parameters are sample-specific and need to be adjusted empirically. To benchmark the automated workflow, we applied on-grid lamella milling with protocols adjusted to five different cell types (Figure 2, Figure 2-figure supplement 1-2, Supplementary table 1). For all samples, lamellae with thickness of roughly 1 μm were generated in 3 steps with FIB currents gradually decreasing from 1 to 0.3 nA as in manual protocols (9, 29). After completion of rough milling for all positions, fine milling to the desired lamella thickness was performed in one or several steps with decreasing FIB currents (Supplementary table 1). Out of a total of 77 positions, we generated 64 lamellae of suitable quality for cryo-ET giving a success rate of 83 % (Table 1, Figure 2-figure supplement 3).

**Table 1:** Statistics of automated on-grid lamellae milling with SerialFIB and corresponding success rates for five different cell types

| Sample | # Target sites | # Polished lamellae | Lamella thickness [nm] |
| --- | --- | --- | --- |
| Sum159 | 22 | 19 | 70-410 |
| HeLa | 22 | 18 | 100-450 |
| *E. huxleyi* | 9 | 9 | 175-470 |
| *C. reinhardtii* | 16 | 10 | 140-350 |
| *S. cerevisiae* | 8 | 8 | 190-300 |
| Total | 77 | 64 |  |
| Success rate [%] |  | 83.1 |  |

### 3D-CLEM targeted lamella preparation

When studying rare biological events, CLEM approaches are indispensable for cryo-FIB-milling, as the final lamella volume captures only a tiny fraction of the entire cell. Using fiducials, e.g. fluorescent microbeads, which are visible in fluorescence, SEM and FIB imaging, cryo-FLM data can be superimposed with the SEM and FIB images to define 3D coordinates of structures of interest and enable targeted lamella preparation. The previously described 3D Correlation Toolbox (3DCT) provides an interface to calculate the geometric transformation between the imaging modalities (30). Here, we introduced new features in 3DCT to improve throughput and usability (Figure 3-figure supplement 1). Using HeLa cells stained while alive with dyes for mitochondria and lipid droplets, we demonstrate the new feature in 3DCT and how SerialFIB interfaces with 3DCT to import correlated positions for automated 3D-CLEM-guided lamella preparation (Figure 3).

**Figure 3:**
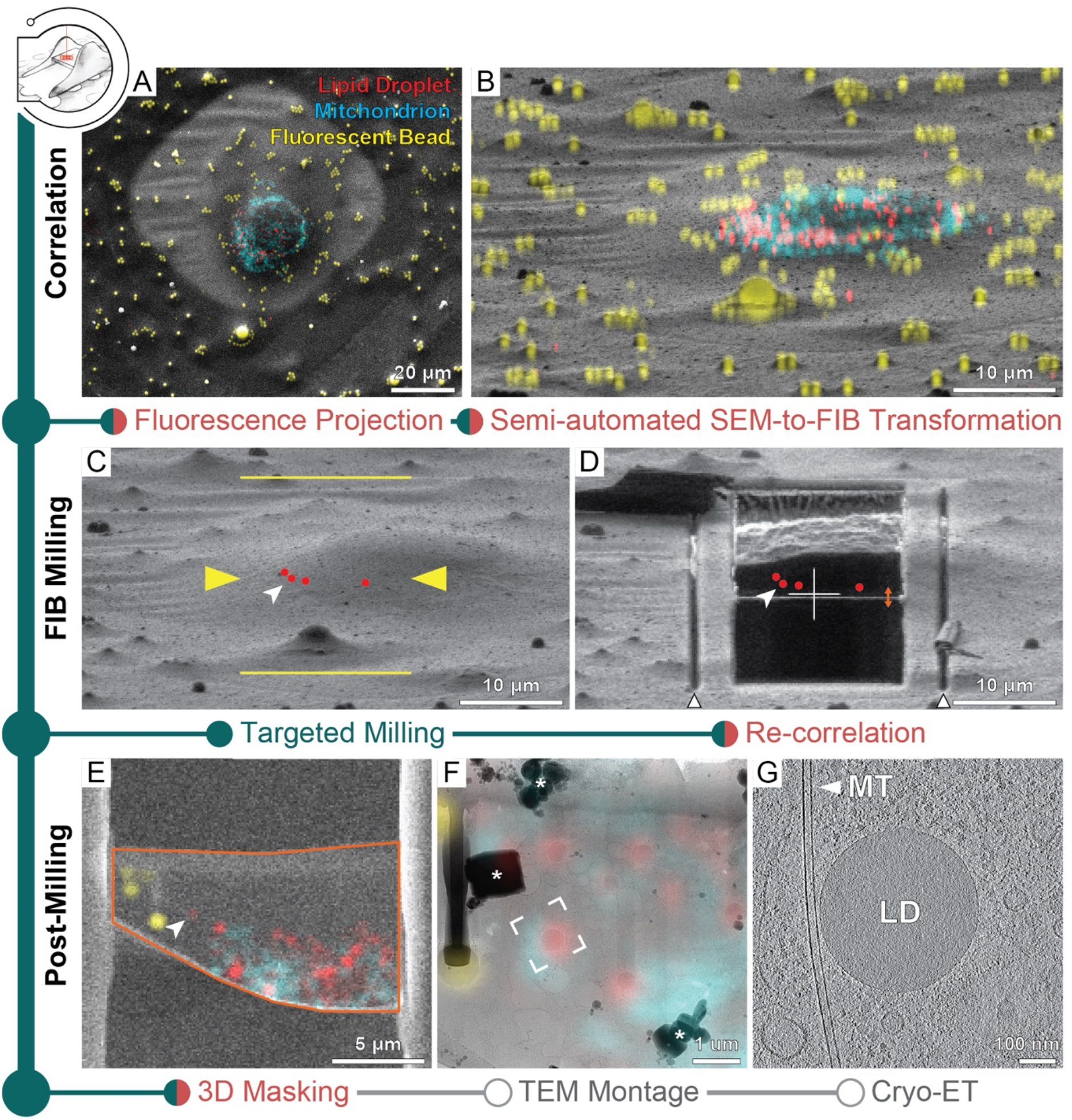
3D-CLEM targeted lamella preparation of HeLa cells. Green modules represent tasks performed with SerialFIB; orange, with 3DCT. (A) SEM and (B) FIB images of a cell. Overlaid are maximum intensity projections of cryo-FLM signal corresponding to lipid droplets red), mitochondria (cyan) and beads (yellow). Correlation with the FIB image is aided by 3D transformation of fiducials detected in the SEM image. (C) Correlated positions of lipid droplets (red dots) imported into SerialFIB. White arrowhead indicates the targeted lipid droplet. Yellow arrowheads indicate the targeted lamella position; yellow lines indicate milling extreme positions. (D) Final lamella after automated FIB milling. Red dots indicate the position of lipid droplets following re-correlation with the FIB image after milling. White triangles highlight micro-expansion joints. (E) Masking of FLM volumes based on lamella outline in the SEM image (orange) and lamella position in the FIB image (orange double arrow in D). A 300 nm FLM slice centered on the lamella is shown. (F) Overlay of the best-fitting fluorescence plane with the TEM image of the lamella. Different heights were sampled in relation to the FIB image to determine the plane where bead and lipid droplet fluorescence signal overlaps best with the TEM image. White frame indicates the targeted lipid droplet and area of acquisition. * denotes ice crystal contamination from transfers. (G) Tomographic slice of area indicated in F depicts the cytosol with a microtubule (MT) and a lipid droplet (LD).

The presence of ice contaminants and the shallow-angle ion beam view of the specimen make the identification of microbead fiducials in a FIB image difficult. Therefore, correlation is often performed first with the SEM image. To aid in this process, we implemented a bead detection tool based on cross-correlation with a Gaussian kernel and Hough transform to facilitate semi-automated detection of beads in the SEM image (Figure 3-figure supplement 1A), after which corresponding beads in the cryo-FLM data can be selected in the 3DCT interface for calculation of a coordinate transform. For fast validation of the calculated transform, maximum intensity projections of the 3D cryo-FLM data can now be generated within 3DCT and overlaid onto the SEM image (Figure 3A). Once the positions of fiducials have been established, their corresponding positions in the FIB image need to be determined. The matching of fiducials between SEM and FIB images was previously performed manually. The new “Fiducial Rotation” feature in 3DCT (Figure 3-figure supplement 1B) takes advantage of the fact that at coincidence height, SEM and FIB images are related by the angle between the two beams and beam shifts. By rotating and translating fiducial coordinates with respect to a user-defined reference point in both images (Figure 3-figure supplement 1B), e.g. surface ice or similarly recognizable feature, the positions of fiducials in the FIB image are determined. These positions can be adjusted subsequently within the 3DCT GUI and a transform calculated. For automated milling, the correlated FIB image is loaded into SerialFIB to serve as the reference image. Maximum intensity projections can be generated and overlaid onto the displayed FIB image to help define milling positions (Figure 3B). Additionally, transformed coordinates of points of interest by 3DCT can be imported into directly into the SerialFIB GUI (Figure 3C). Automated milling as described can then be executed to generate lamellae that retain the structure of interest (Figure 3D).

After milling, 3DCT offers the possibility to overlay the cryo-FLM data with the SEM images of the generated lamella for subsequent navigation during cryo-ET acquisition (Figure 3E-G). A 3D mask describing the thickness, orientation and boundary of the lamella can be created and applied to the original fluorescence volume before projection onto the SEM image (Figure 3-figure supplement 1C). Now implemented in 3DCT, this provides a way to visualize signal contained within the lamella. We term the resulting masked fluorescence volume a FLM “virtual slice” (Figure 3E). For cryo-ET data acquisition, we make use of the 2D affine image registration feature of SerialEM to relate between the SEM/fluorescence overlay and TEM montage of the lamella (Figure 3F). Thereafter, positions are defined for automated tilt series collection (Figure 3F, G).

To evaluate the determinants of successful 3D-CLEM-guided lamella preparation, we targeted lipid droplets in HeLa cells plunge-frozen on two different substrates: titanium grids with holey 1/20 SiO2 support (1 μm holes, 20 μm apart), and the more widely used gold grids with holey 1/4 SiO2 support. Examining the match between FLM virtual slices generated based on correlations with FIB images before milling and the TEM images, we observed that targeted single lipid droplet was retained in 7 out of 10 lamellae on titanium SiO2 1/20 over 3 grids in separate FIB sessions. In contrast, the targeted lipid droplet was retained in 0 out of 4 lamellae on gold SiO2 1/4 from one grid (Supplementary table 2, Figure 3-figure supplement 2). However, upon correlation based on FIB images acquired after milling, none of the correlated target positions coincided with the lamella (Supplementary table 2). To understand the source of the mismatch, we generated virtual slices from the FLM volume at different heights in the FIB view and transformed these slices onto the SEM view and corresponding TEM image. Based on how well bead and lipid droplet fluorescence signals matched the structural features identified in the TEM image, we determined that the best fitting plane was within 3 px distance in y in the FIB image (corresponding to 253–506 nm) of the correlated target position for 9 out of 10 lamellae on titanium SiO2 1/20, and more than 10 px away for all lamellae on gold SiO2 1/4 (Supplementary table 2, Figure 3-figure supplement 2). These data suggest that the milling process may introduce deformations that depend on the local sample properties (26). Elastic 2D registration of FIB images before and after milling revealed a non-uniform displacement in the support of grid squares (Figure 3-figure supplement 3). This displacement occurred irrespective of whether rough milling and polishing were carried out separately, or in one continuous sequence. In all cases, the displacement was more pronounced in areas of thin ice and along the Y direction of the FIB image (up to 2.5 μm), compromising 3D correlation accuracy post-milling.

Our benchmarking results illustrate the importance of having a mechanically stable specimen when performing 3D-CLEM-guided milling for retaining structures of interest in the final lamella. It appears that the combination of a rigid titanium grid, a more continuous support (1/20), and similar thermal expansion behavior between grid and support can help to mitigate local deformations. In cases of mechanical deformations, intracellular fiducials can be leveraged, e.g. by way of lipid droplet staining, to provide local reference points for correlation across all imaging modalities, including TEM.

### Cryo-FIB-SEM volume imaging

Serial cryo-FIB sectioning and SEM imaging produce a 3D nanometer-scale representation of the specimen, providing valuable cellular context on organelle size and distribution (14, 15, 31) and an opportunity to fine-tune the positioning of lamella preparation. This method is performed via iterative ablation of material from a specimen in slices of tens of nanometers followed by SEM imaging of the cross-section. With SerialFIB, we provide an automated cryo-FIB-SEM volume imaging workflow, which can be followed by subsequent lamellae milling. We tested this workflow on Sum159 breast cancer cells. First, an opening into the cell is created by FIB ablation at the cell’s uppermost edge. Then, a volume to be imaged is chosen, within which cellular material is alternatingly ablated in defined steps and the newly exposed surface is imaged by the SEM (Figure 4A, Supplementary video 6). SEM imaging conditions are optimized to visualize cellular structures, while keeping acquisition times as short as possible (Figure 4-figure supplement 1A, see Materials and Methods). As line integration and higher cumulative dwell times are advantageous for noise reduction in cryo-SEM imaging of unstained samples (16), the choice of imaging parameters is a trade-off between acquisition time, image quality and applied dose (between 0.5 – 1.5 e/Å^2^ per slice (32)). For *Chlamydomonas rheinhardtii* cells as an example, improvement of the signal to noise ratio via line integration, with the drawback of longer acquisition times, enabled resolving nuclear pore complexes, the nucleolus and the Golgi apparatus (Figure 4-figure supplement 1E–G).

**Figure 4:**
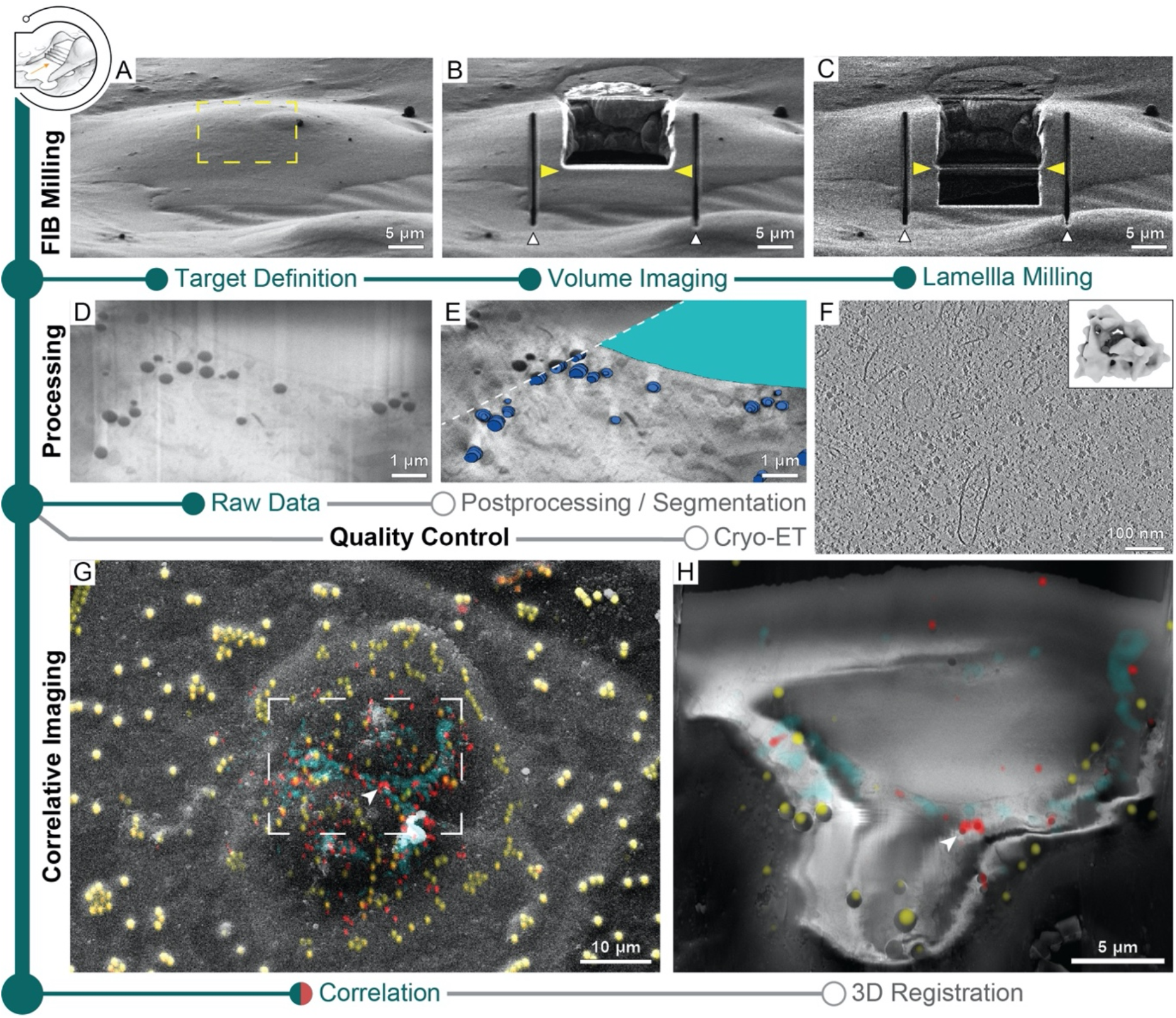
Multi-scale 3D cryogenic imaging of Sum159 breast cancer cells by FIB-SEM volume imaging and cryo-ET. Green modules represent tasks performed with SerialFIB; orange, with 3DCT. Gray modules represent tasks performed externally. (A) FIB image of the cell prior to milling. Yellow rectangle indicates the volume for FIB-SEM volume data acquisition. (B) FIB image of the cell after FIB-SEM volume imaging. Yellow arrows indicate the defined position for lamella preparation. White arrowheads indicate micro-expansion joints prepared after volume imaging. (C) FIB image of the final lamella. (D) Representative slice of the raw FIB-SEM volume data and (D) post-processed data with segmentations of lipid droplets (dark blue) and the nucleus (cyan). (F) Slice through a tomogram acquired from lamella in C. Inset shows the ribosome subtomogram average from data collected on lamellae prepared by this workflow (2 tomograms; 3380 subtomograms; 26 Å resolution). (G) SEM view of two HeLa cells overlaid with a FLM maximum intensity projection of lipid droplet (red), mitochondria (cyan) and fiducial bead (yellow) fluorescence signals, determined by correlation in 3DCT. (H) Representative slice through the post-processed FIB-SEM volume overlaid with a 200-nm virtual slice through the affine-transformed 3D-registered fluorescence volumes.

Subsequently, the stage positions and patterns within the main GUI employed for SEM volume acquisition can be reused to perform lamella preparation (Figure 1-figure supplement 1, stage positions and patterns). Alternatively, the position of lamella generation can be refined manually in the GUI (Figure 1-figure supplement 1, figure 4B, C). In any case, 50-100 nm should be ablated from the last SEM imaging surface to ensure that material potentially damaged by previous imaging and milling is removed. Tomograms acquired from such lamellae are visually indistinguishable from those generated by conventional on-grid approaches (Figure 4F, Supplementary video 7). Ribosome averages from lamellae generated without prior volume imaging reached 24 Å resolution from 4378 subtomograms (Figure 2E inset), while averages from lamellae that were previously volume-imaged reached 26 Å resolution from 3380 subtomograms (Figure 4F inset), indicating that both approaches produce lamellae of similar quality. Streaks in the raw volume data (Figure 4D, Figure 4-figure supplement 1A) resulting from curtaining by dense objects such as lipid droplets can be corrected for using wavelet decomposition (Figure 4-figure supplement 1B) (16). A mask produced by gaussian blurring and image erosion can compensate for charging artifacts (Figure 4-figure supplement 1C). Subsequent local contrast enhancement can restore cellular details (Figure 4E, figure 4-figure supplement 1D). This allowed identification of structures (Figure 4-figure supplement 1E–G) throughout the volume, exemplified by lipid droplets and the nucleus (Figure 4E inlay, Supplementary video 6), which could be further used as internal fiducials for correlation with FLM data.

Using a specimen of HeLa cells stained for mitochondria and lipid droplets, we demonstrate the possibility to combine cryo-FLM with cryo-FIB-SEM volume imaging acquired with SerialFIB. Based on correlation in 3DCT, we defined a window for volume imaging spanning two cells (Figure 4G). Using the centroids of lipid droplets and microbeads in both imaging modalities to fit an affine transform, we were able to register the two volumes (Figure 4H, Figure 4-figure supplement 2). The root-mean-square residual of the fitted transform was 386 nm, with outliers occurring in clusters, indicating potential local elastic image deformations (Figure 4-figure supplement 2, Supplementary video 8). In the present data set, the alignment of SEM slices was also heavily dominated by surface structures outside of the sliced volume. These features presumably lie on a different incline relative to the imaged slices and thus contribute to the shear in Y along Z. Furthermore, stretching of the cryo-FLM volume in Z was necessary, reminiscent of axial distortion correction factors that are applied in light microscopy to account for compression when the refractive index of the immersion medium (air) is lower than that of the specimen (ice) (33). As noted in a recent study where lipid droplet-based registration of cryo-FLM and cryo-FIB-SEM volumes was also performed (31), additional stretching might have been required due to stage drift. Overall, our data indicate that non-affine aberrations in cryo-FLM and SEM imaging, not corrected for here, do not deteriorate the registration accuracy beyond the level required for 3D-CLEM-guided lamella preparation. Thus, such extensions of multi-modal imaging could potentially be employed for advanced preparation of targeted lamellae from voluminous samples.

### Lift-Out lamella preparation

By rapid cooling at elevated pressures, HPF provides a way to preserve fine structure within 200-μm-thick specimens and thus enables the study of multicellular organisms by *in situ* cryo-ET. However, as the resultant specimen is embedded in 50 - 200 μm-thick ice, cryo-FIB lift-out approaches are needed to physically extract material from the bulk specimen for preparation of electron-transparent lamellae.

The time-consuming step in lift-out sample preparations is the generation of trenches that are wide enough to allow access by a gripper-type cryo-micromanipulator and are deep enough such that the lamella captures a meaningful portion of the biological sample. Here, we used SerialFIB for the automated milling of trenches in a high-pressure frozen *Drosophila* egg chamber specimen (Figure 5A–C). We did so by defining a custom pattern sequence file using the Pattern Designer (Figure 1-figure supplement 1C). Due to excessive charging of the HPF sample surface during trench milling, reference image acquisition and all milling steps were performed with “Drift Suppression”, a functionality available through the microscope user interface, whereby the electron beam is used to compensate for positive charges brought in by the ion beam during milling. Trenches were defined in a horseshoe-shaped arrangement to yield a 20 μm x 20 μm extractable chunk from the bulk specimen (Figure 5D). Milling was performed with the specimen surface perpendicular to the ion beam.

**Figure 5:**
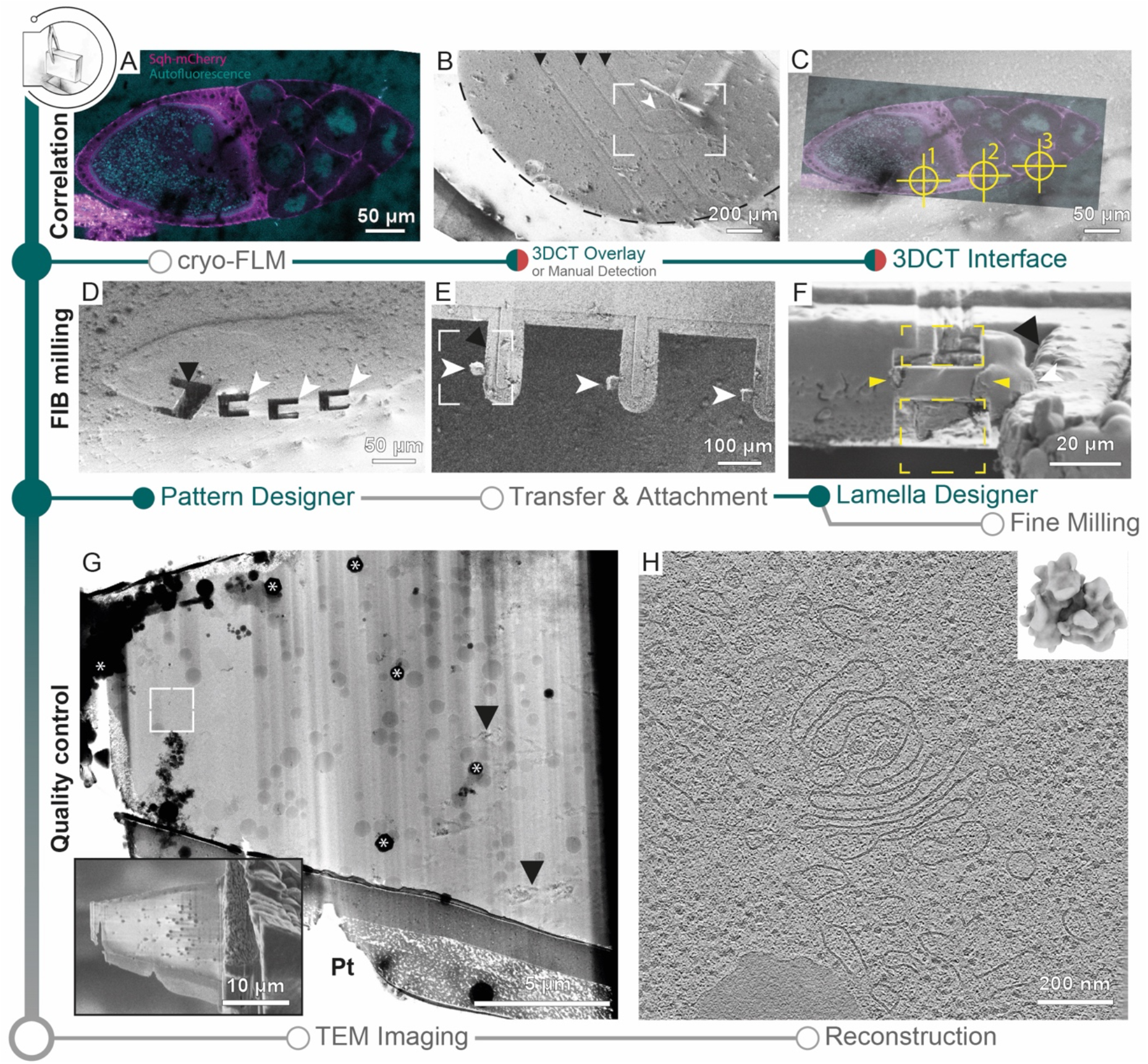
Cryo-FIB lift-out from HPF *D. melanogaster* oocytes. (A) Cryo-FLM image of an oocyte expressing Sqh-mCherry (magenta). Autofluorescence (cyan) indicates oocyte and nurse cell nuclei. The anterior of the egg chamber is to the right. (B) SEM overview of the sample imaged in A. Black arrowheads indicate knife marks introduced during planing in the cryo-microtome. White arrowhead indicates the egg chamber. Dashed line outlines the high pressure freezing planchette. Frame indicates the area depicted in C. (C) Overlay of the cryo-FLM and SEM image of the oocyte. Crosshairs indicate positions chosen for trench preparation, marked 1-3. (D) SEM image of sites prepared for lift-out. Black arrowhead indicates a site produced for FIB-SEM volume imaging to confirm correlation with the fluorescence data, prior to lift-out site preparation (compare Figure 5-figure supplement 1). (E) SEM overview of the half-moon grid after transfer and attachment of the lift-out specimens. White arrowheads indicate the biological material attached to the grid pins (black arrowhead). Frame indicates the region shown in F. (F) FIB image of the lift-out specimen after attachment to a half-moon grid. Yellow arrowheads indicate the milling area for lamella thinning. White arrowhead indicates the lift-out specimen from F. Black arrowhead indicates the edge of the half-moon pin. (G) TEM overview of the lamella from F. Inset shows the SEM image of the final polished lamella prior to transfer to the TEM. White frame denotes the area used for tilt series collection of the tomogram shown in H. Asterisks indicate ice contamination from transfer, arrowheads indicate reflection artefacts from poorly vitrified local regions. Pt: protective platinum layer at the front of the lamella. (H) Single slice through the tomogram collected on the lamella shown in G. Inset shows the *D. melanogaster* ribosome subtomogram average produced from the data set collected on two lift-out lamellae (8 tomograms; 20284 subtomograms).

3DCT can interface with the correlative lift-out module in SerialFIB to identify target positions on the HPF planchette by 2D correlation of cryo-FLM confocal data (Figure 5A) with FIB or SEM images (Figure 5A–C, Figure 5-figure supplement 1). Using the automated procedure, we milled trenches to produce chunks of 20 μm x 20 μm x ~25 μm (Figure 5D). The preparation time of a single position is dependent on the sample and chunk size, and is in the range of 30 - 60 minutes. This adds up to 4 - 5 hours prior to lift-out for five sites. After this step, an undercut is produced by FIB milling at 45° tilt relative to the previous sample orientation leaving the chunk attached to the bulk material on one side only. The prepared site can subsequently be approached by a cryo-gripper mounted on a micromanipulator (11), secured with the gripper arms, detached from the bulk by FIB milling and transferred to a specialized carrier grid, termed half-moon grid (Figure 5E). Unlike previous studies where the half-moon grid was fashioned into slots prior to lamella deposition (11), here we attached the chunks directly to the grid pins. Lamellae attachment was performed by re-deposition of the grid material onto the lamella while held in place by the gripper, similar to procedures described for tungsten needles (22). Subsequently, the gripper was moved to a safe position and organometallic platinum was deposited to secure the lamella to the grid pin (Figure 5E). Finally, the lamellae were polished manually at a pre-tilt of 10° with respect to the grid plane (Figure 5F, Figure 5G inset), and the grid transferred to the TEM for cryo-ET data acquisition (Figure 5G). Tomograms from lamellae prepared with the automated procedures for lift-out site preparation were visually indistinguishable from those prepared via on-grid milling and yield ribosome subtomogram averages of similar quality (Figure 5H).

The presented workflow provides an example of specialized tasks using the pattern sequence file routines for lift-out site preparation based on current state-of-the-art HPF sample carrier, microscope and grid designs. As the technology evolves, new protocols can be designed and implemented through SerialFIB enabling the testing of new geometries and milling strategies to further streamline lift-out procedures.

## Discussion

Here, we present an open-source software tool for the automation of cryo-FIB applications, aimed at improving throughput for *in situ* structural biology. It will be valuable to the growing structural cell biology community in supporting the development of advanced multimodal imaging methods. While communication with the microscope is based on a proprietary API provided by the microscope manufacturer, on-the-fly functions required for automation, e.g. alignment tasks or transformation of FLM data, are decoupled from the commercial scripting interfaces. Therefore, the software architecture allows for flexible development of additional protocols and expansion to other microscope systems by adapting the driver to communication scripts provided by the respective supplier. The modularity of the tools presented here makes it possible to adapt the automation to the specific needs of the biological question at hand.

Optimized cryo-grid preparation remains a stringent requirement for successful automated lamellae micromachining workflows. The distribution of cells on TEM grids is a crucial factor that determines vitrification quality and milling success. To this end, the use of micropatterning to direct the seeding of adherent cells improves their positioning for on-grid lamella preparation (34). Our data and analyses highlight additional factors that must be considered when optimizing grids towards 3D-CLEM-guided FIB preparation:

(i)Mechanical deformation can originate from handling of the specimen, tension within the support due to freezing, or charges introduced by FIB milling or SEM imaging. Two recent hardware solutions have been developed to reduce manual handling of cryo-grids: integration of a fluorescence microscope into the FIB-SEM chamber (35), or the use of shuttles that can be loaded both in the cryo-FLM and cryo-FIB-SEM microscope (22). Tension buildup in the grid support during freezing arises due to differences in thermal expansion coefficients between the metal mesh and the support film (36), or from the tension in the ice film itself (37). In our experiments, titanium grids with almost continuous SiO_2_ support performed better in 3D targeted lamellae preparations than gold SiO2 grids. Nevertheless, local deformations in areas of thin ice were unavoidable during FIB imaging and milling. Thus, grids with stable supports should be considered for cellular preparation and the ice thickness optimized in the vicinity of the milling area.

(ii)Most commercial solutions for diffraction-limited cryo-FLM employ an air objective (14, 38, 39). Thus, refractive index mismatches between the immersion medium (air) and vitreous specimen are unavoidable. This results in a point spread function increasing in asymmetry depending on imaging depth (40), as well as axial compressions of the imaged volume with respect to real object (33). However, for single-cell specimens, where the cellular volumes do not exceed 10 μm thickness, our cryo-FIB-SEM-based analysis of microbeads and lipid droplets in HeLa cells suggests that affine transforms are sufficient to overcome potential aberrations in the imaging and achieving sufficient correlation accuracy for 3D-CLEM-guided milling. Furthermore, native cellular structures such as lipid droplets that can be easily stained with live-dyes provide an interim solution for identifying and validating the cryo-FLM plane corresponding to the produced lamella. This approach would be useful for the targeting and localization of small and rare structures for FIB milling and cryo-ET data collection.

Understanding the caveats of 3D-CLEM at cryogenic conditions through the use case of single-cell samples paves the way for more challenging endeavors, namely identifying and targeting rare biological structures in HPF multicellular specimens. The increasing cooling runtimes of cryo-FIB-SEM microscopes paired with customized automation will enable shifting time-consuming steps of cryo-FIB-SEM volume imaging to overnight operation. While conventional on-grid milling already benefits from the higher throughput made possible by overnight runs and lower in-chamber contamination (41), the impact on successfully targeted cryo-FIB lift-out is expected to be also significant. Currently, cryo-FLM-guided site-specific preparation for *in situ* lift-out relies on manual identification of global features from cryo-FLM data at the surface of the sample for rough registration. Microbead fiducials, if introduced prior to cryo-fixation, would be buried within the bulk specimen and therefore preclude precise registration. Cryo-FIB-SEM volume imaging prior to cryo-FIB lift-out should allow identification of fiducials in the bulk specimen, whether they be fluorescent microspheres or biological landmarks, and subsequent precise registration with the 3D cryo-FLM data. This opens up the possibility of using fluorescence signal to refine targeting for lift-out approaches in the future once on-the-fly post-processing and automated segmentation become available (42–44). In this case, automated overnight operation can increase both throughput and success rate, while the experimenter can focus solely on error-prone steps such as the lift-out procedure or fine milling of the lamellae. The *in situ* lift-out approach is an established technique in material science for room temperature application. With increasing operation experience at cryogenic conditions, further automation will increase throughput of the remaining steps in this workflow, including undercut milling, lamella extraction, transfer and attachment.

The biological research question at hand will dictate the use case. Projects will benefit from automation due to the higher throughput for *in situ* structural biology, improved 3D targeting for on-grid lamella generation, and cutting-edge preparation from voluminous multicellular specimens obtained along with valuable contextual information from FIB-SEM volume imaging. The improved practical application of multimodal imaging under cryogenic conditions through automation holds the potential to accelerate the acquisition of biological insights.

## Acknowledgement

We are grateful to the Walther and Farese labs (Harvard T.H. Chan School of Public Health) for kindly providing Sum159 breast cancer cells, to the Hyman lab (MPI-CBG) for the HeLa cells, Wioleta Dudka for preparation of HeLa cell grids, Zohar Eyal and Assaf Gal (Weizmann Institute of Science) for the *E. huxleyi* grids, and Wojciech Wietrzynski and Ben Engel (Helmholtz Pioneering Campus) for the *C. reinhardtii* grids. We thank members of the Mahamid group, Martin Schorb (Electron Microscopy Core Facility, EMBL), the EMBL cryo-EM platform, Thomas Hoffmann, Florian Beck, Anna Bieber, Cristina Capitanio, and Sagar Khavnekar for invaluable input and support. The cryo-confocal laser scanning microscope (TCS SP8-Cryo CLEM) was developed in collaboration with Leica Microsystems. HKHF was supported by a fellowship from the EMBL Interdisciplinary Postdoctoral Program (EI3POD) under Marie Skłodowska-Curie Actions COFUND (664726). SK was supported by the International Max Planck Research School for Molecular and Cellular Life Sciences. JM acknowledges funding from the EMBL, the European Research Council (ERC 3DCellPhase^-^ 760067) and iNEXT-Discovery project (87103).

## Code and data availability

All code developed in this work is available on GitHub. SerialFIB can be obtained on: https://github.com/sklumpe/SerialFIB. A Python script for wavelet decomposition and post-processing of cryo-FEM-SEM volume imaging data is available in the same repository. The Python 3-ported 3DCT is available on: https://github.com/hermankhfung/3dct. New 3DCT functions described in this work, scripts for cryo-FLM virtual slice series creation, 3D point-based registration and transformation of cryo-FLM data with respect to cryo-FIB-SEM data, and UnwarpJ-based analysis of FIB images before and after milling are available on: https://github.com/hermankhfung/tools3dct.

## Materials and Methods

### Mammalian cell culture and cryo-grid preparation

Hela cells were cultured in DMEM (Thermo Fisher Scientific), supplemented with 10% (v/v) Fetal Bovine Serum (FBS; Biochrom) and 100 mg/mL penicillin-streptomycin (Thermo Fisher Scientific). Sum159 cells were cultured in DMEM/F12, glutaMAX media (Life Technology) supplemented with 5% FBS, 1 μg/ml hydrocortisone (Sigma), 5 ug/ml Insulin (Cell Applications), 10 mM HEPES (pH 7.4), and 100 mg/ml penicillin-streptomycin (Thermo Fisher Scientific). Cells were maintained in TC-25 flasks (Thermo Fisher Scientific) at 37 °C under 5% CO_2_. For EM grid seeding, cells at 60-70% confluence were treated with 0.05% trypsin-EDTA (Fisher Scientific) for 2 min at 37 °C, re-suspended in 1 ml media, passed through a 35 μm mesh Cell Strainer Snap Cap (Corning), and counted. 200,000 cells were seeded into a 35 mm ibidi dish (ibidi) containing up to 8 micropatterned EM grids and 2 mL media, to reach an average surface density of 2×10^4^ cells/cm^2^. Gold or titanium grids, 200 mesh, with 12-nm-thick holey R1/4 and R1.2/20 SiO_2_ film (Quantifoil) were used for the experiments. Grids were passivated, micropatterned and fibronectin-treated according to Toro-Nahuelpan, et al., 2020 (34). Grids were incubated for 2 h with HeLa cells or 1 h with Sum159 cells at 37 °C under 5% CO_2_, and then transferred to a new dish with media for further incubation. Grids were plunge-frozen into liquid ethane at −185 °C 4 - 18 h post-transfer, depending on the desired cell density per square. Plunge-freezing was performed with a Leica GP EM at 37 °C and 75% humidity, and 1 s blot time for all grid types. For correlative experiments, grids were incubated with MitoTracker Green FM (Thermo Fisher Scientific) and BODIPY 558/568 (Thermo Fisher Scientific) at 1:2000 dilutions for 20 min prior to plunge freezing. With the grid mounted on the plunger, 4 μL Crimson Microspheres, 1.0 μm in diameter (Thermo Fisher Scientific), diluted 1:30 in PBS, were applied to the grid from the side containing cells.

### Coccolithophore cryo-grid preparation

*Emiliania huxleyi* strain Eh1 was isolated at the Espegrend Marine Research Field Station, Norway and provided by Prof. Assaf Vardi from the Weizmann Institute of Science. Cells were grown in artificial seawater, supplemented with f/2 nutrient recipe to late exponential phase, under 16 h/8 h light/dark cycles at 18 °C. In order to induce calcification, the external mineral coccolith shell was removed by treating with 20 mM EDTA at pH 8. The de-calcified cells were moved to a calcium depleted medium (100 μM Ca^2+^) for 12 h. Then Ca^2+^ was supplemented to the normal seawater level of 10 mM. This lag time in a calcium depleted medium allowed the cells to resume mineralization in a synchronized fashion upon Ca^2+^ repletion. Cells were centrifuged 3 h after re-calcification at 2000 *g* for 3 min to increase their concentration for plunge freezing. A volume of 4 μL cell suspension at a concentration of 3.07-3.5×10^7^ cells/ml was directly applied to glow-discharged holey carbon R2/1 Cu 200 mesh grids (Quantifoil). After 5 s back-side blotting, plunge-freezing was carried out with a Leica GP EM at 18°C and 90% humidity.

### Budding yeast cryo-grid preparation

The yeast strain used was *Saccharomyces cerevisiae* ΔYAP1801 ΔYAP1802 ΔAPL3 EDE1-eGFP ΔATG15 ΔATG19. Yeast cultures were inoculated from overnight cultures started from colonies, and grown in YPD media at 30 °C to an OD_600_ of 0.8. For grid preparation, 4 μl of cell suspension was applied to holey carbon R2/1 Cu grids (Quantifoil) and plunge-frozen on a Vitrobot Mark IV (Thermo Fisher Scientific) at blot force of 10, blotting time of 10 s, temperature 30 °C and humidity of 90%.

### *Chlamydomonas* cryo-grid preparation

*Chlamydomonas reinhardtii mat3-4* and *CW15* strains were grown with 100 rpm agitation in Tris-Acetate-phosphate Medium (TAP Medium) at room temperature and normal atmosphere under continuous illumination (40 μE white light). Log phase cultures were centrifuged at 2000 *g* for 2 minutes to concentrate the cells 10 times and 4 μl was directly applied to a glow-discharged holey carbon R2/1 Cu 200 mesh grid (Quantifoil). Plunge-freezing was performed using a VitroBot Mark 4 (Thermo Fisher Scientific) with Teflon sheets on both sides and blotting from the back. Blotting time was 10 seconds with a blotting force of 10 at 90% humidity.

### Cryo-FIB lamella preparation

Lamellae were prepared as described in Schaffer et al., 2017 (27) on a dedicated Aquilos dual-beam (FIB-SEM) microscope equipped with a cryo-transfer system and a cryo-stage (Thermo Fisher Scientific). The AutoScript 4 Software is commercially available from Thermo Fisher Scientific.

Cryo-grids with plunge-frozen cells were clipped into autogrids modified with a cut-out for FIB milling, mounted on a shuttle with a 45° pre-tilt (Thermo Fisher Scientific) and transferred into the dual-beam microscope. During FIB operation, samples were kept in high vacuum (< 4 x 10^-7^ mbar) and at constant liquid nitrogen temperature using an open nitrogen-circuit, 360° rotatable cryo-stage. Samples were first coated with organometallic platinum using the gas injection system (GIS, Thermo Fisher Scientific) operated at 28°C, 10.6 mm stage working distance and 8-11 s gas injection time to protect the targeted lamella front during FIB milling. Subsequent sputter coating with platinum (10 mA, 20 s) was performed to improve sample conductivity. Appropriate positions for lamella preparations were identified, registered and the eucentric/coincidence height determined and refined manually or in the MAPS 3.3 software (Thermo Fisher Scientific). Lamellae were prepared at a stage tilt angle of 16-20° using a gallium ion beam at 30 kV. Lamella preparation protocols, including FIB currents and milling times for each cell type are summarized in Supplementary table 1. An example for setting up an automated milling job in SerialFIB including saving positions, defining alignment images and corresponding patterns for each target site is visualized in Supplementary video 1. After polishing, grids were sputter-coated with platinum (10 mA, 5 s), transferred into grid boxes and stored in liquid nitrogen.

### Correlative lamella preparation and analysis

Clipped grids were imaged on a prototype Leica cryo-confocal microscope based on the Leica TCS SP8 system, equipped with a cryo-stage, 50x objective, NA 0.90 and two HyD detectors. Widefield overview montages were acquired in brightfield and fluorescence, and grid squares of interest identified. Z-stacks of HeLa cells stained for mitochondria and lipid droplets and plunge-frozen with fluorescent beads were obtained by 488 nm laser excitation, detecting at 500-545 nm (BODIPY), followed by 552 nm excitation, detecting at 561–630 nm (Mitotracker) and 673-731 nm (microbeads). For setting up of positions in the FIB-SEM microscope, grid squares were identified by overlay of SEM and light microscopy overview images. Coincidence heights were found for each site of interest. SEM and FIB images, encompassing the whole grid square, were acquired for each site. Correlations were established using 3DCT, and the FIB image and 3DCT output were imported into SerialFIB to define lamella positions. Automated rough milling and polishing were performed as above. Lamellae were sputter coated as described. Masked fluorescence projections were generated using 3DCT, overlaid onto the SEM image in Fiji (45), and during a TEM session registered in SerialEM with the lamella TEM montage to guide the setting up of positions for tilt series acquisition. For analysis, registrations between SEM images and TEM montages were refined using BigWarp (46) in Fiji. Fluorescence “virtual slices” corresponding to different heights in the final FIB image were created using the masking utility in 3DCT, projected onto the SEM and the correlated TEM image of lamellae, to identify the fluorescence plane which corresponds best to the structural features in the lamellae (lipid droplets and microbeads). Local mechanical deformation was assessed by elastic registration of FIB images before and after milling using UnwarpJ (47). Broken squares, contaminants, and the site of milling were masked out for this analysis.

### New 3DCT features

(1) For bead detection in the SEM image, pixels are scored based on three measures. First, the image is cross-correlated with a Gaussian kernel, where sigma equals a user-specified bead radius. Second, bead edges are detected with a Canny filter and Hough transform. Pixels more likely to be centers of circles of the given radius are scored higher. The radius is based on the user-provided value and an adjustable multiplier. Third, a soft mask is generated by thresholding of the image with a method of choice, followed by dilation and Gaussian blurring. Scores from all measures are multiplied to obtain a final score. Top peaks are selected and returned as a binary image where peak positions are marked for overlay onto the SEM image in the GUI.
(2) Based on the generated FLM-SEM correlation, fiducial positions in the FIB image are semi-automatically predicted. This is achieved by first calculating a plane that is closest to all fiducials used in the SEM correlation. The user then selects a point in the SEM image on this plane. Rotation around this point by the angle between the electron and ion beam places the fiducials on the FIB image. Finally, fiducial positions are translated according to a matching point in the FIB image selected by the user, to correct for beam shifts and deviation from coincidence height. Fiducial positions are returned in a comma-separated values file that can be imported into the GUI.
(3) To create a 3D mask in the fluorescence volume that matches the position of the generated lamella, a plane is defined by the user by selecting a point in the FIB image and the calculated transform between FLM and FIB images. Given a user-defined polygon in the SEM image, which marks the boundary of the lamella, lines passing through the corner points of this polygon and normal to the SEM image are drawn. The intersection between these lines and plane defined gives the 2D contour of the lamella in 3D space. An additional point placed by the user on the lamella in the SEM image is used to identify the interior of the lamella. A slice of the specified thickness is drawn centered on the contour calculated. The resulting mask can then be applied to the FLM volume during projection generation.
(4) To generate a projection based on a calculated affine transform, FLM volumes are transformed by inverse mapping with linear interpolation, then the maximum intensity projection is taken. This computation is multi-threaded as projections are calculated in patches. To minimize computation, the minimal Z-range that needs to be evaluated to cover the transformed volume is determined before inverse mapping of the FLM data for each patch. Patch size and number of CPUs employed are tuned according to the capacity of the support computer. With this, unbinned fluorescence volumes can be projected onto high-resolution SEM or FIB images during a FIB-SEM session to help guide the setting up of lamella positions in SerialFIB. All features described can be accessed through a PyQt5 graphical interface from command line or from within 3DCT, which has also now been ported to Python 3.

### Cryo-FIB-SEM volume imaging and analysis

For cryo-FIB-SEM volume imaging, samples were prepared as described above. For Sum159 and HeLa cells, SEM images (3072 x 2048, pixel size of 10.377 nm and 19.271 nm) were recorded at 5 kV and 50 pA with a dwell time of 1 μs, using the Everhart-Thornley detector (ETD), with line integration of 16 for noise reduction, resulting in a dose of 0.464 e/Å^2^ per image (32). Milling was performed at 30 kV and 0.5 nA for 10 - 20 s per step using “rectangular cross-section” patterns. Ion beam section thickness was 100 nm, which resulted in step-wise ablation of total volumes of 14.3 μm x 2.0 μm x >13 μm for Sum159, and 25.9 μm x 6.2 μm x >16 μm for HeLa. *Chlamydomonas rheinhardtii* cells were imaged at 3 kV and 13 pA with a dwell time of 0.25 μs, line integration of 64, at a pixel size of 3.4 nm and 6144 x 4096 resolution, corresponding to a dose of 1.12 e/Å^2^ per image, using the ETD, as well as T1 and T2 in-lens detectors. A stage potential was applied in the OptiPlan mode. Milling was performed at 30 kV and 0.3 nA for 60 s with slices of 50 nm.

For volume image analysis, the SEM stack was cropped to the size of the milled area using FIJI (45). Curtaining from ion beam milling was reduced by wavelet decomposition and gaussian blurring of the vertical component (sigma = 6) using pywt (16). Charging was compensated by gaussian blurring (sigma = 35) and subsequent image erosion in two steps to create a mask for image subtraction. The script to perform these tasks is available on the GitHub repository (see Code availability). Image contrast was further enhanced by limited adaptive histogram equalization in FIJI (CLAHE, (48)) with a slope of 3 pixels resulting in restoration of details. Images were aligned using the SIFT algorithm in FIJI (45) and stretched in y by a factor of 1/sin 52° to compensate for foreshortening due to the angle between ion and electron beams. Segmentations were prepared manually in Amira 2020.2 (Thermo Fischer Scientific).

For point-based registration between FIB-SEM and cryo-FLM volumes, centroids of segmented lipid droplets and beads were extracted by connected-component labelling in MATLAB (Mathworks), and by iterative 1D and 2D Gaussian fitting in 3DCT as described (30), respectively. Points were matched in 3DCT using a 2D projection of the segmentation. To avoid local minima, the affine transform relating FLM points to FIB-SEM points was fitted multiple times in Python using (1) the RANSAC approach implemented in OpenCV; (2) L-BFGS-B-based local minimization with different starting positions generated by the TEASER algorithm (49); and (3) basin-hopping coupled with L-BFGS-B-based local minimization. The FLM volumes were transformed according to the affine transform and superposed with the FIB-SEM volume in Amira for visualization. Python scripts for the affine registration and volume transformation are available on GitHub (see Code availability).

### Fly husbandry

*Drosophila melanogaster w[*]; P{w[+mC]=sqh-mCherry.M}3* (FlyBase ID: FBst0059024 (50)), expressing an mCherry-tagged Sqh protein (myosin regulatory light chain) under the control of the *sqh* endogenous promoter, were obtained from the Bloomington *Drosophila* Stock Center (BL-59024) and maintained at 22 °C on standard cornmeal agar. Flies were transferred into a fresh vial supplemented with yeast paste 24 h prior to dissection of egg chambers for HPF.

### High pressure freezing and cryo-FIB lift-out

*Drosophila melanogaster* egg chambers were dissected from ovaries in Schneider’s medium and high pressure frozen (HPM010, Abrafluid) in the 100 μm cavity of gold-coated copper Type A carriers (Engineering office M. Wohlwend) using Schneider’s medium containing 20% Ficoll (70 kDa) as filling medium. HPF carriers were soaked in hexadecene and blotted on filter paper prior to freezing. Cryo-planing of approximately 40 μm of the surface was performed in a cryo-microtome (EM UC6/FC6 cryo-microtome, Leica Microsystems) using a 45° diamond trimming knife with a clearance angle of 6°C (DiAtome). The sample was glued to a custom 3D-printed shuttle after the Leica design (22) using cryo-glue (2:3 ethanol:isopropanol mixture) for cryo-confocal imaging with a Leica TCS SP8 equipped with Leica EM cryo-CLEM stage. Imaging was performed using a HC PL APO 50x/0.9 DRY objective in fluorescence mode using a 552 nm excitation laser at 17% total laser strength and 488 nm excitation at 6.8% total laser strength, corresponding to roughly 0.94 mW and 0.38 mW, respectively (based on measured laser intensity values). Detection channels were 650-751 for mCherry and 495-545 nm for autofluorescence using HyD detectors at a pinhole size of 2 Airy units. Z-stacks of 38 μm from the surface were collected using a step size of 1 μm. Fluorescence signal decreased significantly with imaging depth. Targeting of positions for automated trench milling was based on the surface topology of the sample produced by the cryo-planing. Positions were chosen based on registration with cryo-FLM data in 3DCT, which allowed the identification of regions within the egg chamber.

For lift-out experiments, an Aquilos dual-beam FIB-SEM microscope (Thermo Fisher Scientific) was equipped with a Kleindiek MM3A-EM micromanipulator and a cryo-gripper head (Kleindiek Nanotechnik) cooled by copper wires attached to the microscope anti-contaminator. Standard 45° pre-tilt shuttles were used for accommodating both HPF carrier and Auto-Grid clipped receiver grid on an Aquilos stage that was modified by removing ~2 mm of material of the stage using a file to allow the cryo-gripper to reach the sample surface. OmniProbe four-post Cu half-grids were used as receiver grids (Plano EM). After specimen loading, platinum GIS deposition was performed for 10 seconds at 27 °C with the GIS needle distance at a stage working distance of 10.6 mm. Sample were sputter coated with platinum in the chamber (10 mA, 15 s). For all trench milling steps and SerialFIB session setup, drift-suppression using the SEM was applied to compensate positive charges brought in by FIB milling. The stage was rotated by 180° relative to loading position and tilted to 7° to orient the sample surface perpendicular to the FIB. Trench milling was performed with “regular cross-section patterns” at 30 kV, 1 nA creating a C-shape using SerialFIB (Figure 5). The milling included two patterns of 40 μm x 15 μm symmetrically arranged around a 20 μm block. Together with a third pattern, 10 μm x 40 μm generated at an offset of 15 μm to the left and perpendicular to the previous two, yielding a 20 μm x 20 μm block for lift-out. Milling time per position was 30 minutes. In total, 5 positions were prepared, resulting in a milling preparation time of 2.5 h. Undercuts were performed manually by tilting the stage to 43°C and employing “cleaning cross-section” patterns of 1 μm height, 22 μm width and 15 μm depth at ~7 μm distance from the sample surface. Lift-out was performed with the microgripper oriented parallel to the FIB angle and perpendicular to the sample surface. The produced sample chunks were approached by the gripper. After contact of both tweezer arms with the sample, the chunk was milled loose from the bulk material with a “regular cross-section patterns” at 1 nA with a pattern of 20 μm in length, 1 μm in width until extraction from the bulk was possible. Subsequently, the microgripper was raised until it was ~400 μm above the sample surface. The stage was moved down by ~15 mm into safe distance from the gripper and subsequently moved to the receiver grid position while keeping the Z-height locked. Then, the stage was lifted to the appropriate Z-height of the receiver grid. The microgripper was moved to the post of the TEM half-grid until achieving contact of the sample. “Regular cross-section” patterns at height of 2 μm and width of 6 μm were used to attach the sample to the post by re-deposition of the post material onto the sample. Subsequently, the gripper was opened, releasing the chunk that has been welded to the pin, and moved to a safe position. This is achieved by lifting the gripper until it is 400 μm above the sample surface and then moving the stage down by 15 mm, making enough room to safely move it to the park position. This procedure can be performed for all four pins. Finally, deposition of platinum GIS for 3 x 30 s in 3-min intervals at a stage working distance of 9 mm was used to strengthen the sample attachment to the pin. Lamella production was done manually with sequentially reduced FIB currents of 3 nA to 5 μm lamella thickness, 1 nA to 4 μm, 0.5 nA to 3 μm, 0.3 nA to 2 μm, and 0.1 nA to 1 μm using “regular cross-sections” patterns. Subsequent polishing was performed using “regular patterns” at 0.1 nA to 600 nm and 50 pA to 200 nm. The FIB divergence was compensated by over- and under-tilts of 1° (27). The lamella was milled until contrast started fading in SEM imaging at 3 kV, 13 pA.

### Cryo-electron Tomography

Cryo-ET data was acquired on a Titan Krios (Thermo Fisher Scientific) at 300 kV, equipped with a K2 Summit direct detection camera (Gatan) operating in dose fractionation mode and a Quantum post-column energy filter (Gatan). Autogrids containing lamellae were loaded such that the axis of the pre-tilt introduced by FIB milling was aligned perpendicular to the tilt axis of the microscope. At an EFTEM magnification of 42000 and a resulting pixel size of 3.37 Å or 3.52 Å, up to 10 tilt series were collected on a single lamella in low dose mode using SerialEM (24). Starting from the lamella pre-tilt, images were acquired at 2.5 - 4.5 μm underfocus, in 2° increments between +/-62° using a dose-symmetric tilt scheme (51). The total dose was around 130 e^-^/Å^2^ with a constant electron dose per tilt image. Tilt series were either collected with a 70 μm objective aperture or a Volta phase plate (VPP, Thermo Fisher Scientific) with prior conditioning for 6 min.

### Tomogram reconstruction

Data were preprocessed in Warp using movie frame alignment to compensate for beam-induced movement, CTF estimation and tilt stack sorting (52) or TOMOMAN, kindly provided by W. Wan, personal communication, 2021, employing MotionCorr2 (53), NovaCTF (54) and CTFFIND4 (55). In etomo (IMOD software package (56)), four times binned tilt images (movie sums) were aligned using patch tracking and tomograms reconstructed via weighted back projection.

### Subtomogram analysis

Coordinates of ribosomes in Sum159 cells were determined in 4x binned tomograms by template matching utilizing the pyTOM toolbox (57), using a down sampled human 60S large ribosomal subunit (Gaussian filter with sigma 2, EMDB-2938) as a template and a spherical mask with 337 Å diameter. After manual selection, 4378 and 3380 ribosomal particles were localized in tomograms from standard on-grid FIB-milled lamellae (4 tomograms) and lamellae prepared after FIB-SEM volume imaging (2 tomograms), respectively. Subtomograms and corresponding CTF models were reconstructed in Warp with a box size of 140 px, a pixel size of 3.37 Å, and assuming a 350 Å particle diameter for normalization. Subtomogram alignment and averaging was performed in RELION (version 3.0.7, (58)). Particles from both datasets were pooled to generate an initial average via 3D classification into one class with the human 80S ribosome (EMDB-2938), low-pass-filtered to 60 Å, as reference. The resulting average was 3D-refined. Then, particles were re-grouped based on their origin, i.e. from standard FIB-milled lamellae or from lamellae prepared after FIB-SEM volume imaging. Both particle groups were refined separately starting from the alignments of the pooled average and its density as reference. Resolutions of the two final reconstructions were calculated via Fourier shell correlation of two half-maps, and the reconstructions were filtered in the RELION post-processing step to their respective resolutions of 24 Å and 26 Å (0.143 FSC) for particles from tomograms of standard on-grid FIB-milled lamellae and lamellae prepared after FIB-SEM volume imaging, respectively.

*Drosophila* ribosomal particle positions were determined in 4x binned tomograms by template matching in STOPGAP using a down-sampled reference from a previously determined structure of the *Drosophila melanogaster* ribosome (EMD-5591). In total, 20284 ribosomal particles from 8 tomograms were picked. Subsequent subtomogram alignment and averaging was performed in STOPGAP (59). Particles were extracted with a box size of 64 px, at a pixel size of 7.04 Å. The resolution was calculated via Fourier shell cross-correlation of half-maps to be 24.0 Å (0.5 FSC) and 20.8 Å (0.143 FSC).

Subtomogram averages were visualized with the UCSF ChimeraX package (60).

### Python packages

Implementations were done in python3. Packages used in this work were numpy, scipy, scikit-image, open-cv, tiffile, psutil, PyQt5, pickle and matplotlib.

