## Supplementary file for "A Modular Platform for Streamlining Automated Cryo-FIB Workflows"

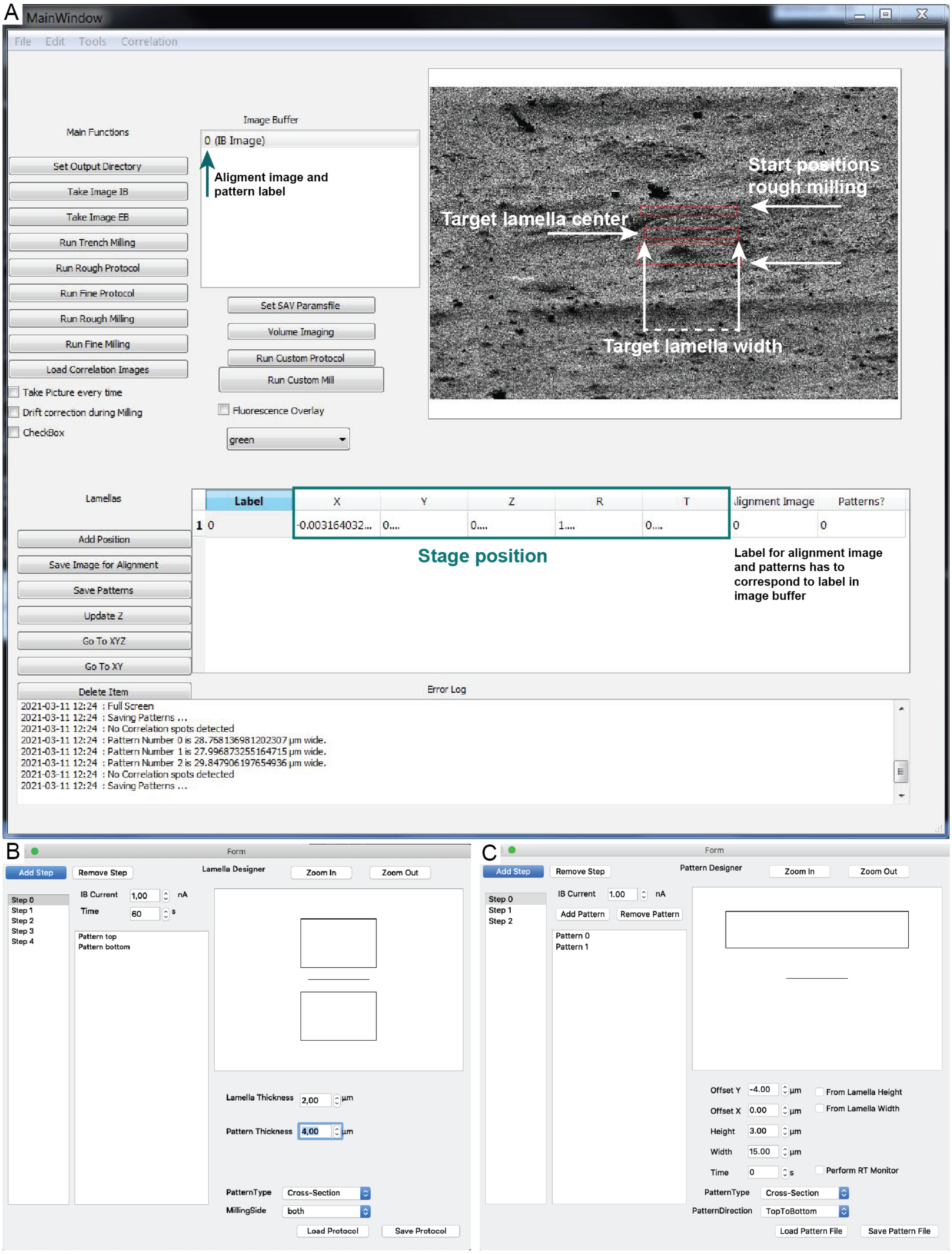


**Figure 1-figure supplement 1:** Graphical user-interface (GUI) for automated cryo-FIB protocols. (A) The GUI displays buttons for the main functions (imaging and several FIB milling procedures; left), the image buffer where reference images are selected and displayed in the alignment window to prepare and adjust milling patterns (top right, red rectangles). Specialized applications (volume imaging, custom milling procedures and fluorescence overlay) are available below the image buffer.

In the coordinate navigator (bottom), milling sites are connected with alignment images and corresponding patterns. From the Tools tab, Lamella Designer, Pattern Designer, Volume Designer and Script Editor can be selected. (B) The Lamella Designer allows for stepwise construction of user-defined patterns and milling procedures. (C) The Pattern Designer enables creation of customized patterns, e.g. for lift-out trenches.

**Supplementary video 1:** Setting up automated on-grid lamella milling in SerialFIB.

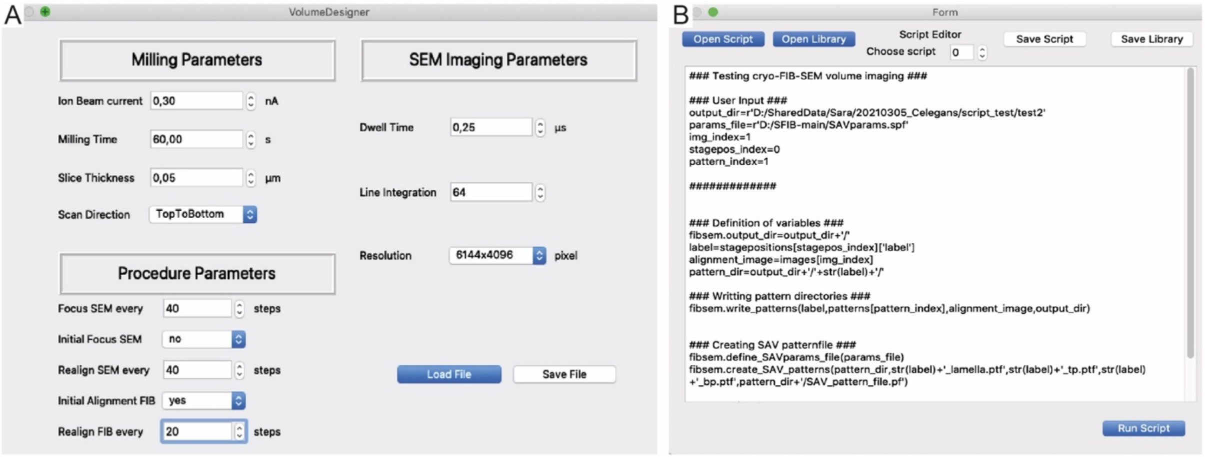


**Figure 1-figure supplement 2:** SerialFIB interface for volume imaging. (A) In the Volume Designer, FIB-SEM volume imaging workflows can be constructed with user-definable FIB slicing and SEM imaging parameters. (B) The Script Editor offers a scriptable interface for customized milling protocols which can be saved as independent scripts or within libraries.

**Supplementary video 2:** Tomographic volume of a Sum159 breast cancer cell related to figure 2E depicting the cytosol with microtubules (MT) and a mitochondrion (Mito) next to the nucleus.

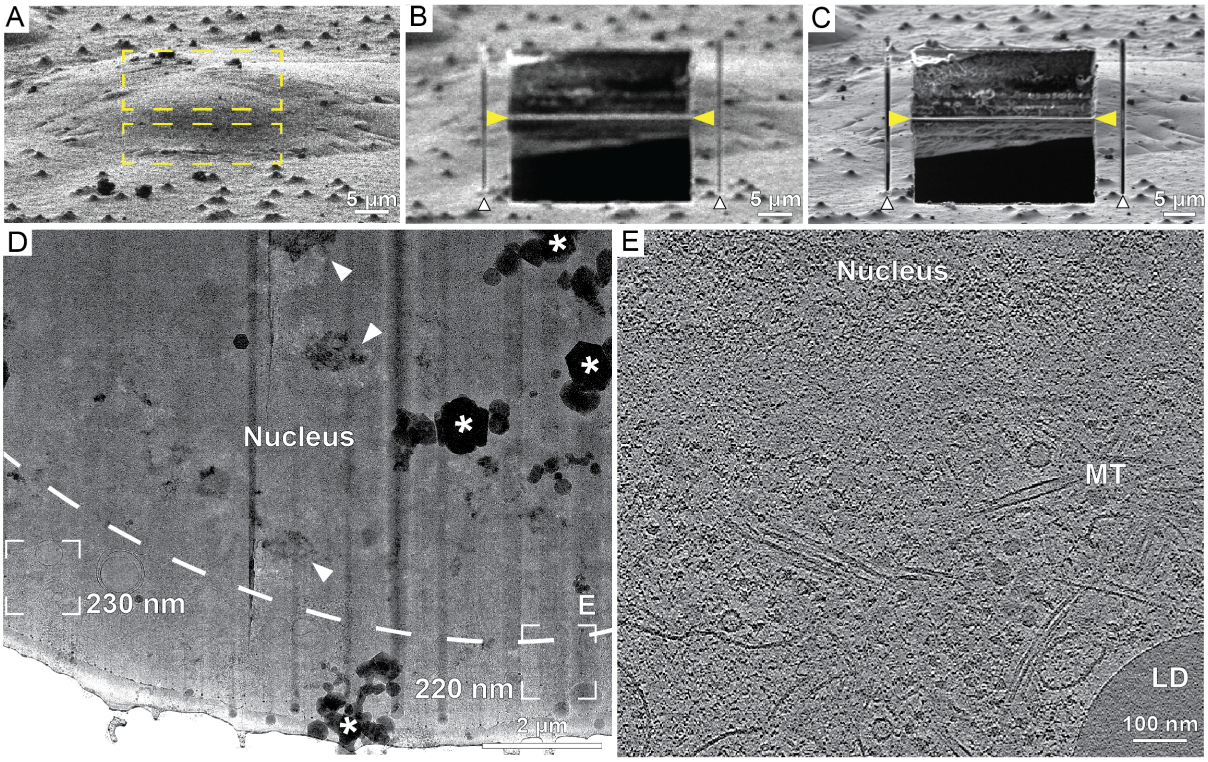


**Figure 2-figure supplement 1:** Automated on-grid lamella preparation of HeLa cells. (A) FIB image of a cell prior to lamella preparation. Yellow rectangles indicate milling patterns where material is subsequently removed. (B) Micro-expansion joints (white arrowheads) and rough lamella (yellow arrowheads) milled to a target thickness of 1 µm. (C) Lamella after polishing to a target thickness of 300 nm. (D) TEM overview of lamella in C. Frames indicate examples of tilt-series acquisition positions (out of 3 acquired on this lamella) and the local thickness determined from reconstructed tomograms. Nuclear envelope delineated by a dashed line. Vertical dark stripes (curtains) commonly originate from dense lipid droplets, that resist milling. Milling protocol was adjusted to minimize curtaining (Table S1). * denotes ice crystal contamination from transfer between FIB and TEM. Arrowheads point to crystalline ice reflections in the nucleus. Area indicated by E is enlarged. (E) A slice through the tomogram depicts the cytosol with microtubules (MT), a lipid droplet (LD) and the nucleus.

**Supplementary video 3:** Tomographic volume of a HeLa cell related to figure 2-figure supplement 1E depicting the nuclear periphery, and the cytosol with microtubules (MT) and a lipid droplet (LD).

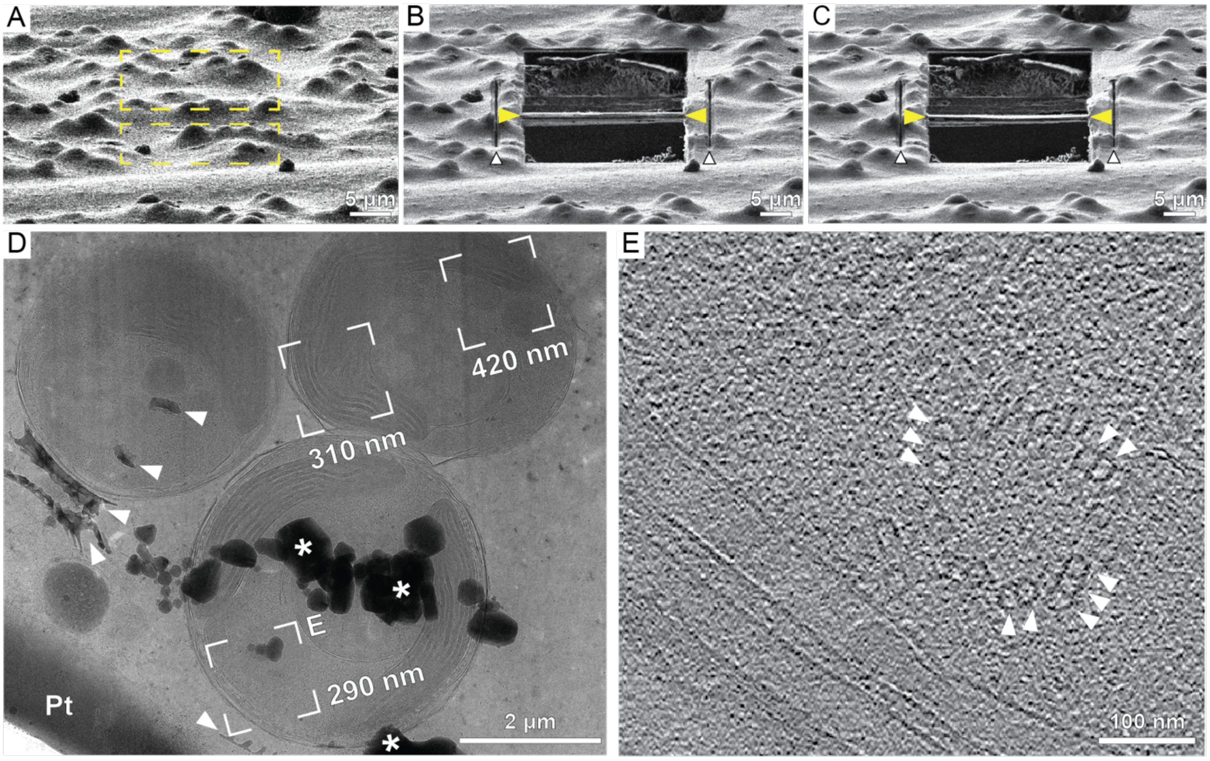


**Figure 2-figure supplement 2:** Automated on-grid lamella preparation of *E. huxleyi* cells. (A) FIB image of an agglomeration of cells prior to lamella preparation. Yellow rectangles indicate milling patterns where material is subsequently removed. (B) Micro-expansion joints (white arrowheads) and rough lamella (yellow arrowheads) milled to a thickness smaller than the target of 1 µm due to prolonged milling times. (C) Final lamella after polishing to a target thickness of 300 nm. (D) Section of the TEM overview of the lamella showing three individual cells. Frames indicate examples of tilt-series acquisition positions (out of 10 acquired on this lamella) and the local thickness determined from reconstructed tomograms. Dense structures like intra- and extracellular calcium carbonate crystals (top two and bottom three white arrowheads, respectively) are observable, as well as the protective Pt layer. Milling protocol was adjusted to minimize curtaining from the dense crystalline structures (Table S1). * denotes ice crystal contamination from transfer between FIB and TEM. (E) A slice through the tomogram indicated by E in the TEM overview in D depicts the cytosol with microtubules (white arrowheads) assembled into doublets and triplets constituting a basal body of a cilium.

**Supplementary video 4:** Tomographic volume of *E. huxleyi* cells related to figure 2-figure supplement 2E depicting the cytosol with the basal body of a cilium.

**Supplementary table 1:** Selected parameters for milling of micro-expansion joints, lamella milling and polishing of five different cell types

| Sample | Micro-expansion joint milling | | | | Rough milling | | | | Polishing | | | |
|---|---|---|---|---|---|---|---|---|---|---|---|---|
| | distance from lamella [μm] | width [μm] | current [nA] | time [sec] | step | nominal lamella thickness [μm] | current [nA] | time [sec] | step | nominal lamella thickness [nm] | current [pA] | time [sec] |
| Sum159 | 4 | 0.3 | 1 | 30 | 1 | 5 | 1 | 480 | 1 | 400 | 100 | 210 |
| | | | | | 2 | 3 | 0.5 | 210 | 2 | 300 | 50 | 150 |
| | | | | | 3 | 1 | 0.3 | 210 | | | | |
| HeLa | 4 | 0.3 | 1 | 30 | 1 | 5 | 1 | 540 | 1 | 300 | 50 | 360 |
| | | | | | 2 | 3 | 0.5 | 300 | | | | |
| | | | | | 3 | 1 | 0.3 | 270 | | | | |
| *E. huxleyi* | 4 | 0.3 | 1 | 120 | 1 | 5 | 1 | 600 | 1 | 800 | 100 | 240 |
| | | | | | 2 | 3 | 0.5 | 480 | 2 | 600 | 50 | 240 |
| | | | | | 3 | 1 | 0.3 | 480 | 3 | 300 | 30 | 240 |
| *C. rheinhardtii* | 5 | 0.5 | 0.3 | 60 | 1 | 5 | 0.3 | 210 | 1 | 800 | 50 | 180 |
| | | | | | 2 | 3 | 0.3 | 120 | 2 | 600 | 50 | 90 |
| | | | | | 3 | 1 | 0.1 | 120 | 3 | 400 | 50 | 60 |
| *S. cerevisiae* | 5 | 0.5 | 0.3 | 60 | 1 | 5 | 0.3 | 210 | 1 | 800 | 50 | 180 |
| | | | | | 2 | 3 | 0.3 | 120 | 2 | 600 | 50 | 90 |
| | | | | | 3 | 1 | 0.1 | 120 | 3 | 400 | 50 | 60 |

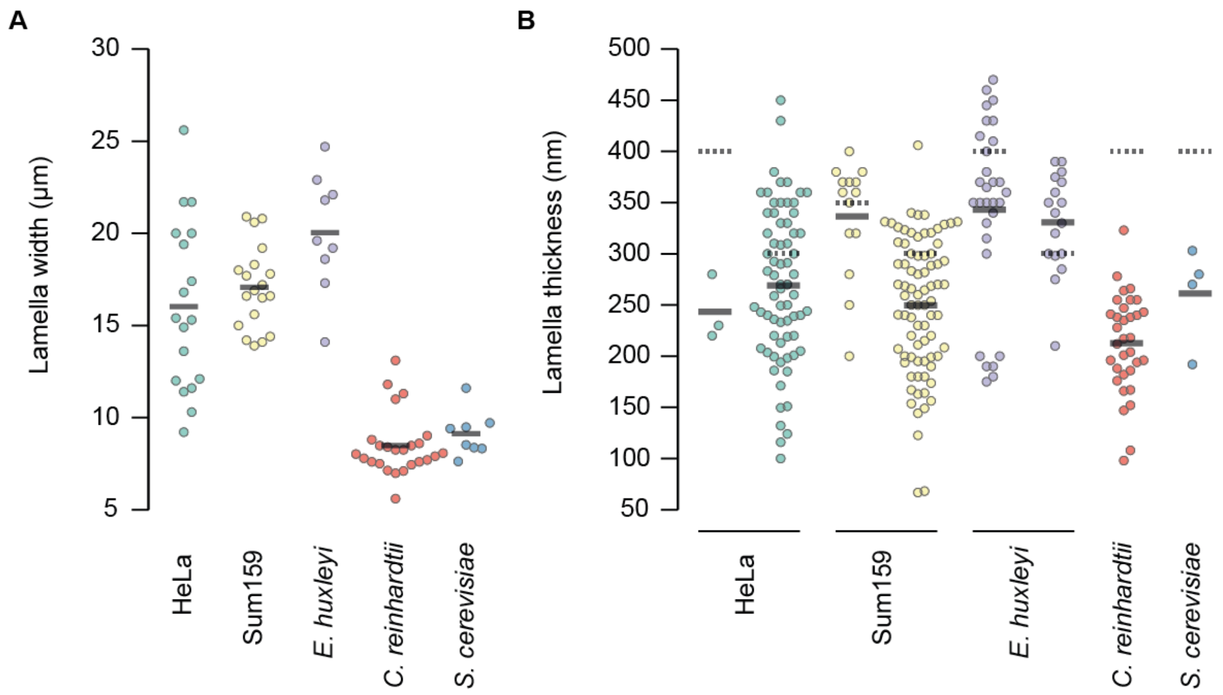


**Figure 2-figure supplement 3:** Width (A) and thickness distribution (B) of successfully prepared lamellae from four eukaryotic cell types. Thickness measurements were made per tomogram acquired. Solid line indicates the mean thickness; dotted line indicates the target thickness.

**Supplementary video 5:** Tomographic volume of a HeLa cell related to figure 3G depicting the cytosol with microtubules (MT), a lipid droplet (LD) and a multi vesicular body (MVB). * denotes ice reflections originating from incomplete vitrifcation.

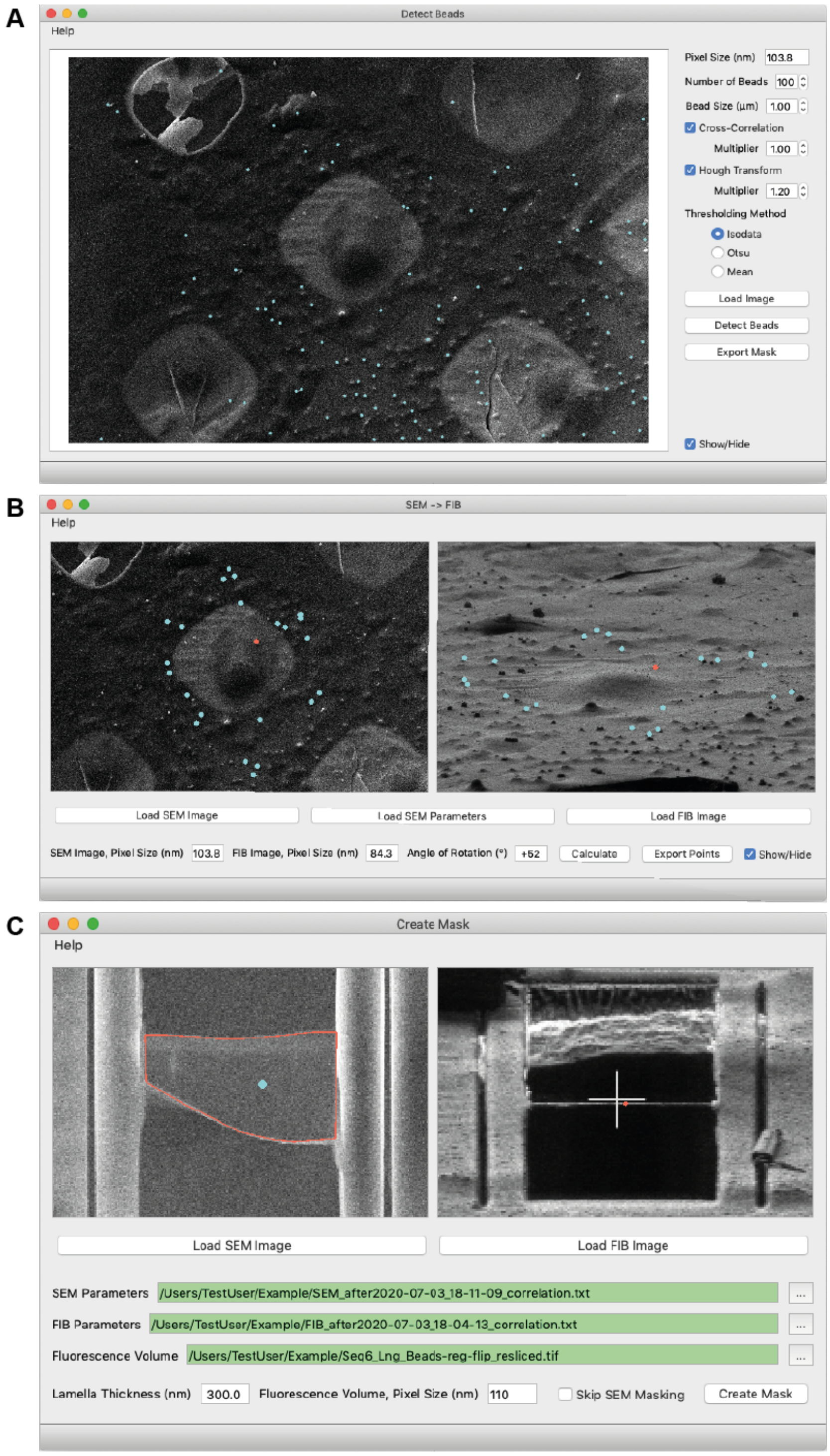


**Figure 3-figure supplement 1:** New 3DCT features. (A) Semi-automated detection of beads in SEM images. Different methods for detection are available. Blue dots indicate beads found. (B) Rotation of fiducials from SEM (left) to FIB (right) orientation. Blue dots in the SEM image indicate fiducial beads used for correlation. Red dot in both images indicates a common feature identified visually, which serves as the center of rotation. Blue dots in the FIB image indicate the rotated positions. (C) 3D mask creation for generation of a virtual slice in the cryo-FLM volume. Red outline in the SEM image (left) indicates lamella boundary, and blue dot indicates lamella interior. Red dot in the FIB image (right) indicates the lamella position.

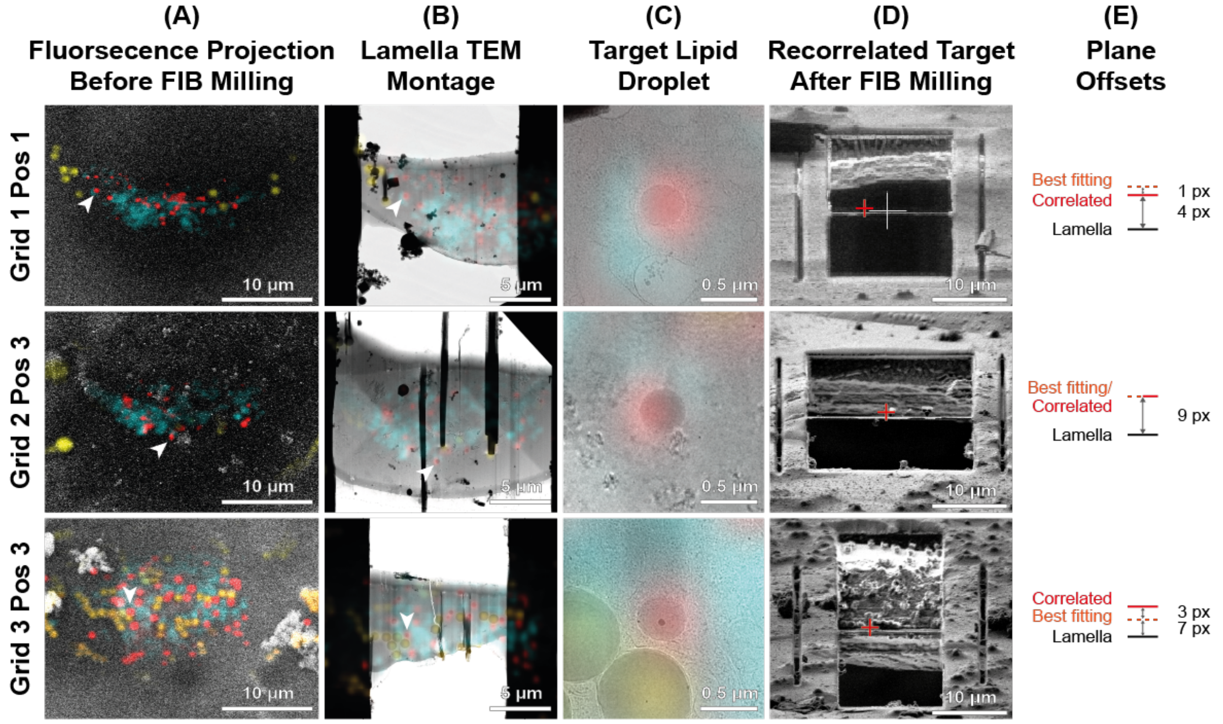


**Figure 3-figure supplement 2:** Examples of successful 3D-targeted milling of lipid droplet. (A) Cryo-FLM volumes projected onto SEM images before FIB milling. Red, lipid droplet; cyan, mitochondria; yellow, fiducial beads. White arrowhead indicates the targeted lipid droplet. (B) The best-fitting fluorescence plane projected onto the TEM montage of the lamella, determined by sampling of different heights in the FIB image using the 3DCT masking utility (see Material and Methods). (C) Zoom-in view of the targeted lipid droplet overlaid with the best-fitting fluorescence plane. (D) FIB view of the final lamella. Red cross indicates the correlated position of the target lipid droplet based on the final FIB image. (E) Offsets of correlated positions post-milling and positions of the best fitting plane relative to the final lamella position in the FIB image, tabulated in Supplementary table 2.

**Supplementary table 2:** Positions of correlated lipid droplet and best fitting fluorescence plane on lamellae.

| Grid | Grid type | FIB image pixel size (nm) | Grid position | Targeted lipid droplet visible in TEM? ‡ | Correlated Y position relative to lamella (px) | Best fitting plane relative to correlated position (px) |
|---|---|---|---|---|---|---|
| 1 | Ti $SiO_2$ 1/20 | 168.7 | 1 | yes | 4 | 1 |
| | | | 2 | yes | 3 | 3 |
| | | | 3 | yes | 11 | -2 |
| | | | 4 | yes | 8 | -1 |
| 2 | Ti $SiO_2$ 1/20 | 84.3 | 1 | yes | 2 | 3 |
| | | | 2 | no | 7 | 2 |
| | | | 3 | yes | 9 | 0 |
| 3 | Ti $SiO_2$ 1/20 | 84.3 | 1 | no | 9 | 3 |
| | | | 2 | no | 14 | -3 |
| | | | 3 | yes | 7 | -3 |
| 4 | Au $SiO_2$ 1/4 | 67.5 | 1 | N/A | -1 | N/A |
| | | | 2 | no | -2 | 11 |
| | | | 3 | no | -9 | 22 |
| | | | 4 | no | -8 | 19 |

‡ N/A = assessment could not be made due to severe condensation on lamella

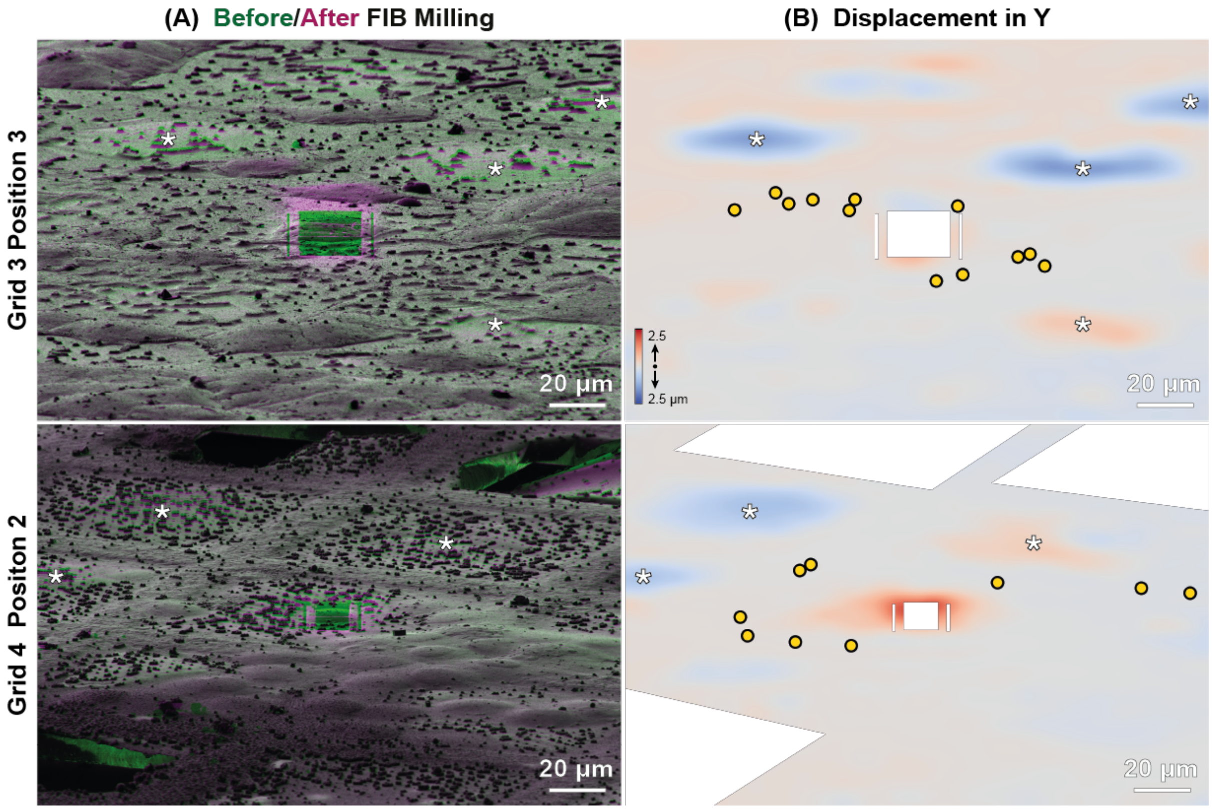


**Figure 3-figure supplement 3:** Mechanical deformations of specimens. (A) Overlay of exemplary FIB images of HeLa cells on Ti $SiO_2$ 1/20 (top) or Au $SiO_2$ 1/4 (bottom), before (green) and after milling (magenta) (related to Supplementary table 2). (B) Heat map depicting displacements in Y as determined by elastic registration. Red, up; blue, down. Yellow circles represent beads used for correlation in 3DCT. White areas were excluded from the analysis, which included broken squares, surface contaminants and the site of milling. White asterisks indicate unmilled squares which exhibited strong deformations (up to 2.5 µm).

**Supplementary video 6:** Cryo-FIB-SEM volume of a Sum159 cell depicting raw and post-processed data with segmentations of lipid droplets (blue) and the nucleus (cyan).

**Supplementary video 7:** Tomographic volume of a Sum159 breast cancer cell related to figure 4F depicting the cytosol with two vault structures and a vesicle (V).

**Supplementary video 8:** FIB-SEM volume of HeLa cells in Figure 4H, superposed with lipid droplet and bead segmentations and fluorescence volumes transformed.

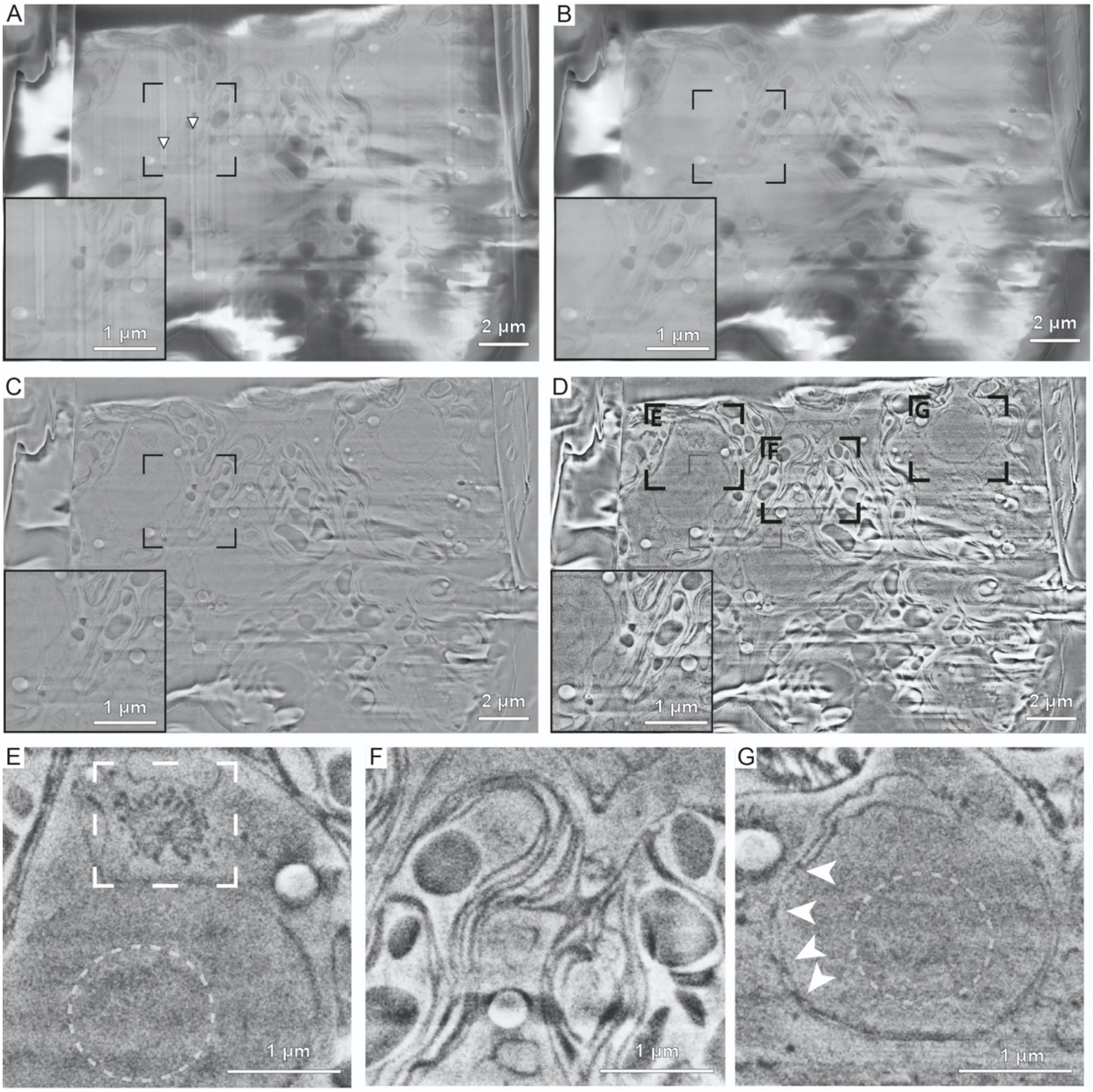


**Figure 4-figure supplement 1:** Serial FIB volume imaging and processing of *Chlamydomonas reinhardtii* and subsequent image processing adapted from (1). (A) Raw data. Arrowheads indicate curtaining artefacts from ion beam milling. (B) De-striped image using wavelet decomposition to 8th level, Coiflets family 3 and gaussian blurring (sigma 6) of the vertical component. (C) Local charge adjusted by image subtraction of a gaussian blurred (sigma = 35) and subsequently three times eroded image from the image in B. (D) Image after median filtering (3.0 pixels) and local contrast enhancement using CLAHE (slope 3.0). (E-G) Zoom into regions indicated in D, showing a Golgi stack (E, rectangle), thylakoid membranes (F), the nucleolus (E and G, dashed circles) and nuclear pore complexes in the nuclear envelope (G, arrowheads).

A

| Rotation | | Scale | | Shear | | | | Translation |
|---|---|---|---|---|---|---|---|---|
| $\varphi$ | 25.6° | $X$ | 5.409 | $s_{xy}$ | -0.029 | $s_{xz}$ | -0.061 | $x$ -695.2 px |
| $\theta$ | 9.8° | $Y$ | 5.486 | $s_{yx}$ | -0.032 | $s_{yz}$ | 0.869 | $y$ -5668.3 px |
| $\psi$ | 0.0° | $Z$ | 6.259 | $s_{zx}$ | -0.008 | $s_{zy}$ | -0.003 | $z$ -1297.4 px |

B

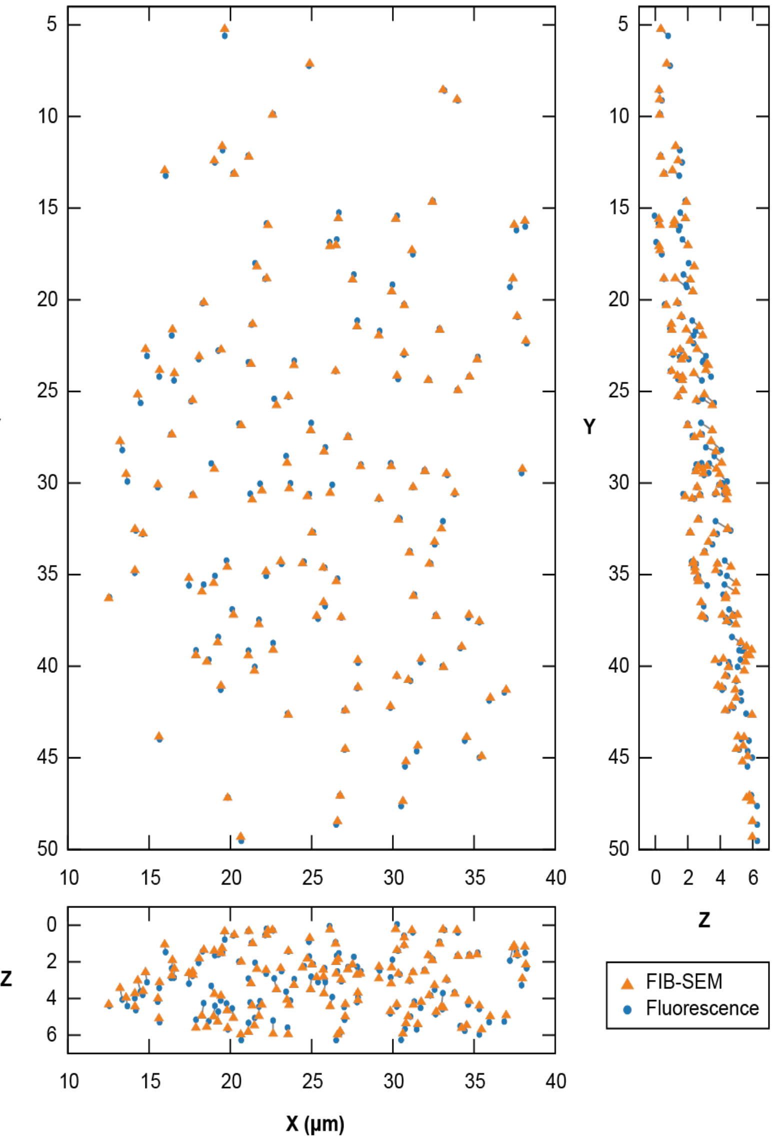


**Figure 4-figure supplement 2:** Point-based affine 3D registration between FIB-SEM and fluorescence volumes. Registration was based on the unweighted centroids of lipid droplets and beads segmented in the FIB-SEM data, and centers of their fluorescence signal from Gaussian fitting in the corresponding FLM volume ($n$ = 133). Lipid droplet and bead coordinates were pooled and used equally for the registration. (A) Fitted parameters for transformation of the FLM volume. Euler rotation angles are given in ZXZ convention. Shear factor $s_{yz}$ represents shearing in Y along the Z axis. (B) 2D orthogonal projections of the FIB-SEM and transformed fluorescence coordinates. A gray line connects matching points between the two volumes. The root-mean-square-deviation of the fit was 386 nm, with residuals ranging from 29 nm to 882 nm.

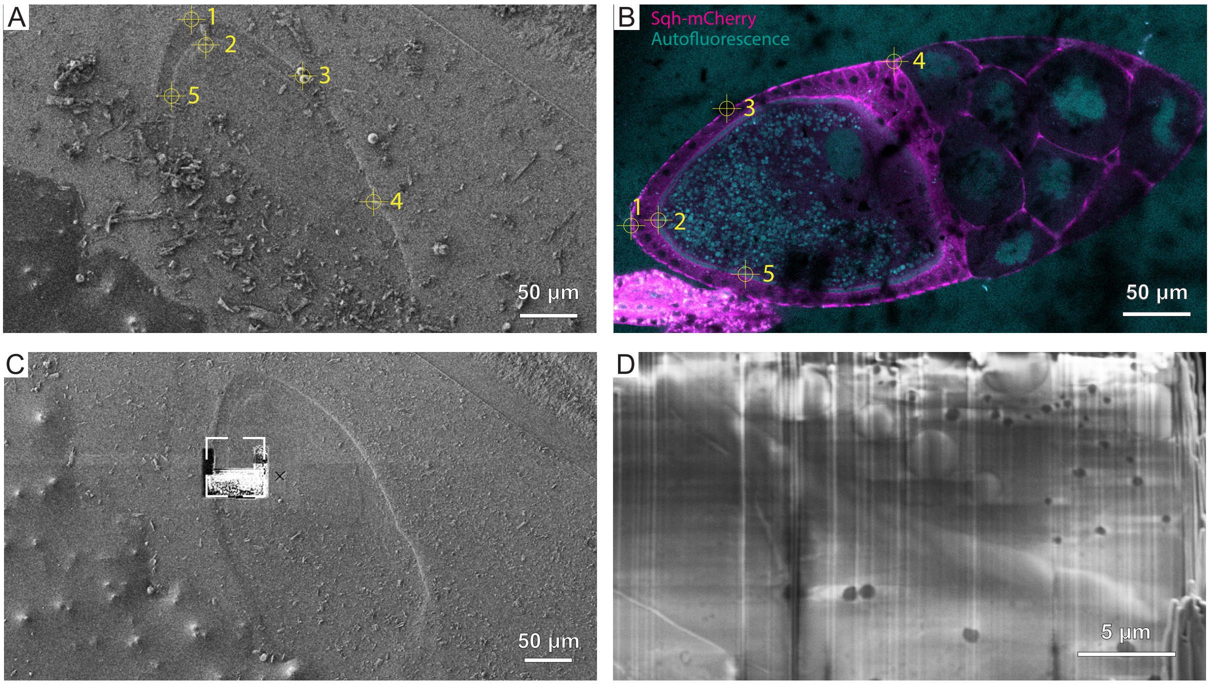


**Figure 5-figure supplement 1:** Correlation for cryo-FIB lift-out from HPF *D. melanogaster* oocyte. (A) FIB and (B) cryo-FLM image (anterior to the right) used for correlation. Crosshair and numbering indicate features used for registration of the two imaging modalities in 3DCT. (C) FIB image of the trench milled for SEM imaging. Frame indicates the region imaged in D. (D) SEM image of the prepared surface.

**Supplementary video 9:** Tomographic volume from a *D. melanogaster* liftout lamellae related to figure 5H depicting the cytosol with a lipid droplet (LD) and a Golgi apparatus.
